# Efficient uniform sampling explains non-uniform memory of narrative stories

**DOI:** 10.1101/2025.07.31.667952

**Authors:** Jianing Mu, Alison R. Preston, Alexander G. Huth

## Abstract

Humans do not remember all experiences uniformly. We remember certain moments better than others, and central gist better than detail. Current theories focus exclusively on surprise to explain why some moments are better remembered, and do not explain gist memory. We propose that humans uniformly sample incoming information in time, which explains both non-uniform memory and gist. Rather than surprise, this model predicts that the mutual information between a given moment and the rest of the experience drives memory. To test this model, participants listened to narrative stories and recalled them immediately afterward. Using large language models to quantify the information structure of narrative stories and participants’ recall, we found that our parsimonious uniform sampling model explained memory better than earlier theories. These findings suggest an alternative, simpler account of human memory that does not rely on costly feedback mechanisms for prioritizing encoding of specific information.

## 1 Introduction

We experience a constant stream of information every day, and yet we do not remember everything equally well [1, 2]. For example, why are some parts of narrative stories remembered better than others [3, 4]? One leading theory holds that surprising moments lead to the subjective perception of event boundaries and hence are better remembered [5, 6]. However, surprisal poorly predicts event boundaries in narratives [7], questioning the validity of this account. Selectively encoding event boundaries also requires constant, resource-intensive top-down control. And because this strategy focuses on selectively encoding specific moments, it fails to explain how humans extract the global gist of a narrative, a crucial ability for understanding continuous experience. Instead, an efficient memory system must consider that natural experiences do not unfold randomly. Narratives are highly structured because they are governed by the physical properties of the world and the communicator’s intent [8, 9, 10]. This structure allows humans to learn patterns that they can use to compress information and predict future experiences [11, 12, 13]. We propose that taking advantage of these patterns obviates the need to selectively encode stimulus information entirely. Memory behaviors explained by selective encoding, such as better memory at event boundaries, can emerge from a simple mechanism in which humans uniformly sample incoming information from a structured stimulus. We show that uniform sampling at a constant rate better accounts for memory than selective encoding and also explains individuals’ ability to recall global narratives.

To assess how humans use the structure of our experience to form memories, one major challenge is quantifying the patterns in naturalistic stimuli. Past research has used easily quantifiable, discrete stimuli to show that humans efficiently perceive [14, 15, 16] and remember [17, 18, 19, 20, 21] information. However, naturalistic stimuli such as narratives have more complex statistics. We overcome this challenge by using large language models (LLMs) to estimate the information structure of narrative stories. These powerful models were trained to capture patterns in natural language based on massive text corpora. We used LLMs to measure mutual information between different moments of a narrative stimulus (Fig. 1a), as well as individuals’ recall of these moments (Fig. 1b). These mutual information estimates enable us to measure how much information is shared across moments of a narrative (green outlines in Fig. 1a, overlapping areas in Fig. 1c). For example, in the first part of the story the narrator mentions, “I quit the job”, which shares mutual information with a later statement, “I’m gonna find my next path”. This significantly extends current models of memory of narratives, because models that only explain memory for discrete moments could not account for the semantic and causal relationships across parts of the story [22, 23].

**Figure 1.**
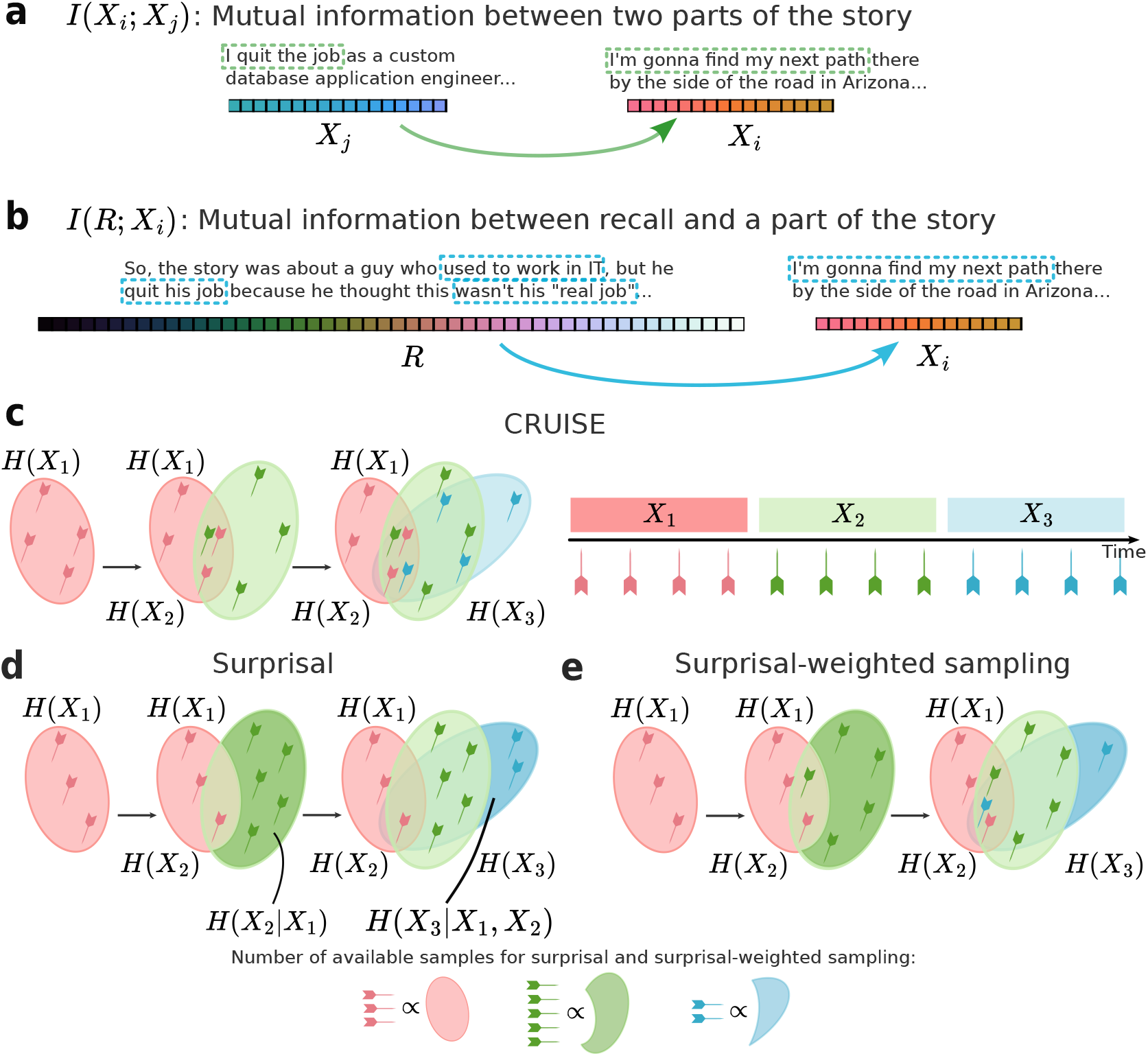
Leveraging mutual information estimates to assess memory. **a.** Mutual information between different parts of a story shows how much is shared. **b**. Mutual information between part of a story and a person’s recall of the story shows how much is remembered. **c**. Our proposed model, constant-rate uniform incremental sampling for encoding (CRUISE). Ovals represent the information in each part (*X*_1_, *X*_2_, *X*_3_) of a story. The listener samples the information during each part uniformly in time (darts). Because information is shared across the story (overlapping areas), samples for one part can also pertain to other parts. The amount remembered about each part is the total number of samples that fall within it, meaning that parts with more shared information will be remembered better. **d**. Event segmentation theory proposes that humans selectively encode surprising information. Here, samples for each part of the story only fall within the surprising information of that part (dark shaded regions). The number of samples taken during each part of the story is proportional to surprisal. **e**. A middle ground, surprisal-weighted sampling, combines features of CRUISE and the surprisal model. It is similar to CRUISE in that one still samples uniformly within the total information of each part of the story, but here the number of samples is proportional to a part’s surprisal instead of duration, as in CRUISE.

The ability to calculate mutual information across a narrative enables us to make two important predictions about memory under our uniform sampling model. First, uniform sampling prioritizes information shared across the story, unlike selective encoding models that prioritize surprise. If each part of the story is uniformly sampled, parts that share more information with other parts (overlapping colors in Fig. 1c) will effectively be sampled more frequently. We predict that those parts will be better remembered. Second, by oversampling information shared across many parts of a story, uniform sampling also automatically prioritizes extracting the central gist of a narrative. The level of gist versus detail in memory should then correspond to the overall sampling rate during encoding. Selective encoding, by focusing on unique information, fails to account for the extraction of gist.

To test these predictions, we recruited participants to listen to eight narrative stories, mark event boundaries, and then immediately recall the stories. The uniform sampling model better predicts participant recall data than selective encoding. Quantifying information characteristics also showed that effects usually attributed to event boundaries are equally well explained by uniform sampling. Further, uniform sampling explains gist extraction, as central information shared across an entire narrative is automatically oversampled. Overall, these results suggest that human memory of natural narratives is best explained by simple uniform sampling of incoming information.

## 2 Results

### 2.1 How do we sample information to encode into memory?

To understand how humans remember natural narratives, we consider the scenario in which a human participant *Y* listens to a story *X*, and then recalls the story out loud, producing a recall *R*_*Y*_. We first divide the story into *n* sequential parts *X*_1_, *X*_2_,…, *X*_*n*_, in which a “part” is an arbitrary span of time in seconds. If the listener uniformly samples information at a constant rate, then the number of bits sampled during each part of the story should be the duration of that part multiplied by the sampling rate. Because the story is a natural narrative, each part will share some information with the other parts. Thus, after the listener has heard the entire story, the total amount of information accumulated about part *X*_*i*_ will be larger than just the amount sampled during *X*_*i*_, as some of the information sampled during other parts also pertains to *X*_*i*_ (Fig. 1c). The amount of information gained about *X*_*i*_ during another part *X*_*j*_ will be proportional to the amount of information shared between *X*_*i*_ and *X*_*j*_ (overlapping areas in Fig. 1c) divided by the total information in *X*_*j*_ (ovals in Fig. 1c), measured here in bits. The fraction of *X*_*j*_ bits that pertain to *X*_*i*_ can then be multiplied by the duration of *X*_*j*_ and sampling rate to find the number of bits about *X*_*i*_ that were sampled during *X*_*j*_. Finally, the total number of bits accumulated for *X*_*i*_ across the story will be the sum of these values across all parts.

We assume that the number of bits listener *Y* remembers about part *X*_*i*_, i.e., the mutual information between *X*_*i*_ and their recall *R*_*Y*_, is proportional to the number of bits that the listener accumulated about *X*_*i*_. Because both this “memory rate” and the sampling rate are unknown constants, we treat their product as a single sampling×memory rate constant, *g*. We instantiate this proposed model mathematically as the Constant-Rate Uniform Incremental Sampling for Encoding (CRUISE) model, in which the amount of information that a listener will remember about the *i*-th part of the story, *I*(*X*_*i*_; *R*), is

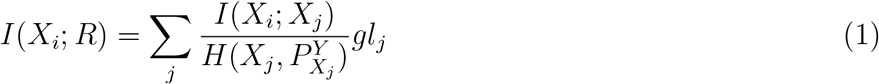

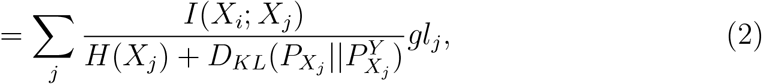

where *I*(*X*_*i*_; *X*_*j*_) is the mutual information between parts *X*_*i*_ and *X*_*j*_ measured in bits; *H*(*X*_*j*_) is the total information in part *X*_*j*_ in bits (i.e., the amount of information in *X*_*j*_ when it is presented out of context); *g* is the rate of sampling memory in bits per second; *l*_*j*_ is the length of *X*_*j*_ in seconds; and 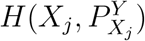 is the cross-entropy of *X*_*j*_ under listener *Y* ‘s model of the world, indicating the total number of bits listener *Y* needs to perfectly recall *X*_*j*_. By the definition of cross entropy, 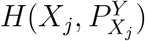 is the sum of the true entropy *H*(*X*_*j*_) and a term that captures how different the listener’s model of the world 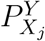 is from the true distribution underlying the story 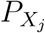, i.e., the Kullback–Leibler divergence of 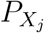 from 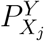 (See details in Supplemental Methods: Assumptions of CRUISE). In Fig. 1c, *I*(*X*_*i*_; *R*) is the total number of samples falling within *H*(*X*_*i*_), and *I*(*X*_*i*_; *X*_*j*_) is the overlapping area between *H*(*X*_*i*_) and *H*(*X*_*j*_).

Because we cannot explicitly measure how well a listener’s world model captures the statistics of the stimuli, we assume that the cost of representing *X*_*j*_ under listener *Y* ‘s model is proportional to the amount of information in *X*_*j*_,

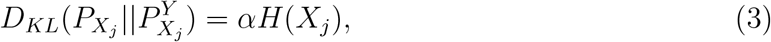

where *α* is a positive, unitless constant that represents how inefficiently the listener models this story. Thus, CRUISE (Eq. 2) becomes

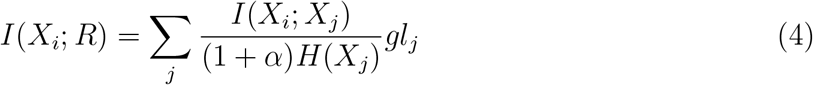

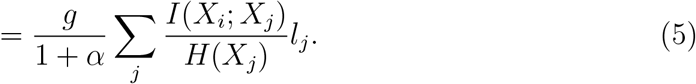

This form conveniently combines the unknown factors *g* and *α* into a single multiplicative constant 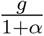 that will be estimated using linear regression.

An alternative proposition to CRUISE is that humans selectively encode surprising information [5]. Under this theory, the number of bits a listener remembers about a part of the story should be proportional to its surprisal (Fig. 1d):

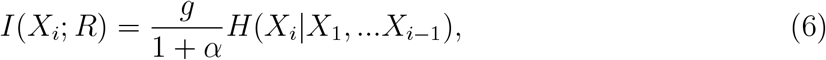

in which *g* is a unitless sampling rate in bits stored per bit perceived and *H*(*X*_*i*_ |*X*_1_, …*X*_*i*−1_) is the surprisal of part *X*_*i*_. In Fig. 1d, the samples only fall into the shaded regions *H*(*X*_*i*_ |*X*_1_, …*X*_*i*−1_), and the number of samples is proportional to the area of *H*(*X*_*i*_ |*X*_1_, …*X*_*i*−1_). We have expressed this theory in similar terms to CRUISE to facilitate comparison. Notably, because this theory only focuses on surprisal, it does not consider shared information.

We also consider a third formulation, which we call surprisal-weighted sampling, that combines CRUISE with selective encoding. Under this model, information is still sampled uniformly within each part of the story, but the number of bits sampled is proportional to surprisal rather than duration (Fig. 1e). This differs from the surprisal model in that it factors in the shared information structure in the narrative, and differs from the CRUISE model in that it preferentially samples surprising parts of the story. This model is given by

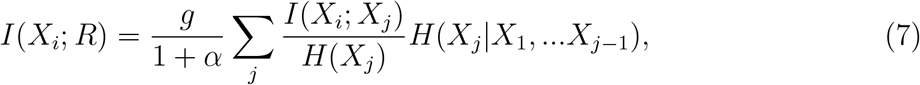

where the sampling rate *g* is unitless (bits per bit) and tells us what fraction of the surprising bits lead to samples. Note that if information density is perfectly uniform across time [24, 25, 26, 27, 28], then *H*(*X*_*j*_ |*X*_1_, …*X*_*j−*1_) ∝ *l*_*j*_ and surprisal-weighted sampling is identical to CRUISE. However, if information density varies, CRUISE and surprisal-weighted sampling will make different predictions.

### 2.2 Estimating information quantities

To evaluate these models we used LLMs to estimate the entropy of text, something that was previously intractable using traditional experimental methods. At each position in text, LLMs output a probability distribution over possible next tokens—roughly equivalent but not identical to words. This distribution is used to calculate cross entropy, which is the amount of information added by each token that could not be predicted from the preceding tokens in the story. The total information is thus the sum of cross entropy across all tokens. If two parts of a story share information, the second part will have lower cross entropy if the LLM has already read the first part, as each token will be predicted more accurately using the shared information. This reduction in cross entropy defines mutual information between the two parts (*I*(*X*_*i*_; *X*_*j*_) = *H*(*X*_*j*_) − *H*(*X*_*j*_ |*X*_*i*_); see details in Methods: Information estimates). We use the same method to estimate the mutual information between the recall and each part of the story, *I*(*X*_*i*_; *R*). This metric closely tracks human annotations of recall (Fig. S3).

To validate LLMs’ ability to capture mutual information we performed several analyses. We know that mutual information should be symmetric, i.e., *I*(*A*; *B*) = *I*(*B*; *A*), and non-negative, i.e., *I*(*A*; *B*) ≥ 0. We used these theoretical properties to evaluate the reliability of six open LLMs and selected the LLM with the best symmetry and non-negativity scores among those with the needed capabilities, Llama3-8b-Instruct [29], for further analyses (see Methods: Estimating the mutual information of recall and a part of the story *I*(*R*; *X*_*i*_)). Experimental results were robust to choice of LLM (see Fig. S4).

### 2.3 Experiment and model evaluation

To test CRUISE against selective encoding, we performed a behavioral experiment in which participants listened to narrative stories from *The Moth Radio Hour* while simultaneously performing event segmentation (pressing a button whenever, in their judgment, one meaningful event ends and the next one begins [30]), and then verbally recalled the stories immediately afterward. These event segmentation judgments allowed us to later inspect participants’ non-uniform memory at consensus event boundaries. We used eight narrative stories, each of which was heard and recalled by about 50 participants recruited from Prolific, yielding a total of 413 participants (216 female, 184 male, 13 non-binary, see Table S1). The mean story word count was 1834 words (mean duration: 11.19 minutes) and the mean recall word count was 429 words. We transcribed the recalls using Whisper-large-v3 and manually checked the automated transcripts for errors (see Methods: Behavioral data analysis).

Using this dataset we tested how well three models—CRUISE, surprisal, and surprisal-weighted sampling—predicted participants’ recall behavior. We first divided each story into non-overlapping windows of approximately equal duration, *X*_1_,…, *X*_*n*_, with the beginning and end of each window adjusted minimally to match phrase boundaries (see Methods: Creating equal-duration windows of story text). We measured *I*(*X*_*i*_; *R*) for each participant and then averaged across participants, giving a single value showing how well each window was remembered on average. We then fit linear regression models to predict the mean *I*(*X*_*i*_; *R*) across participants using each of the CRUISE, selective encoding, and surprisal-weighted sampling models. Each regression model included the constant term 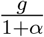, which is estimated as a regression coefficient. Initial examination of the data showed that participants tended to produce recalls of similar duration regardless of the story duration, leading to lower apparent sampling rate for longer stories. Our experiment required recalls to reach a minimum length but did not give any incentive for longer recalls, so this likely reflects a reward-maximizing strategy for the participants. To account for this effect we fit a separate slope term for each story.

Results from these regression models show that both CRUISE and surprisal-weighted sampling predicted the mean amount of information participants recall about each window quite well (CRUISE: *Adj*.*R*^2^ = 0.385, surprisal-weighted sampling: *Adj*.*R*^2^ = 0.322, Fig. 2a,b). In contrast, the surprisal-only model was a poor predictor of recall (Fig. 2c, *Adj*.*R*^2^ = 0.151). The effectiveness of CRUISE and surprisal-weighted sampling was not explained by the general tendency of some text to share more information with others (See Supplemental Results: Ruling out the shared information confound).

**Figure 2.**
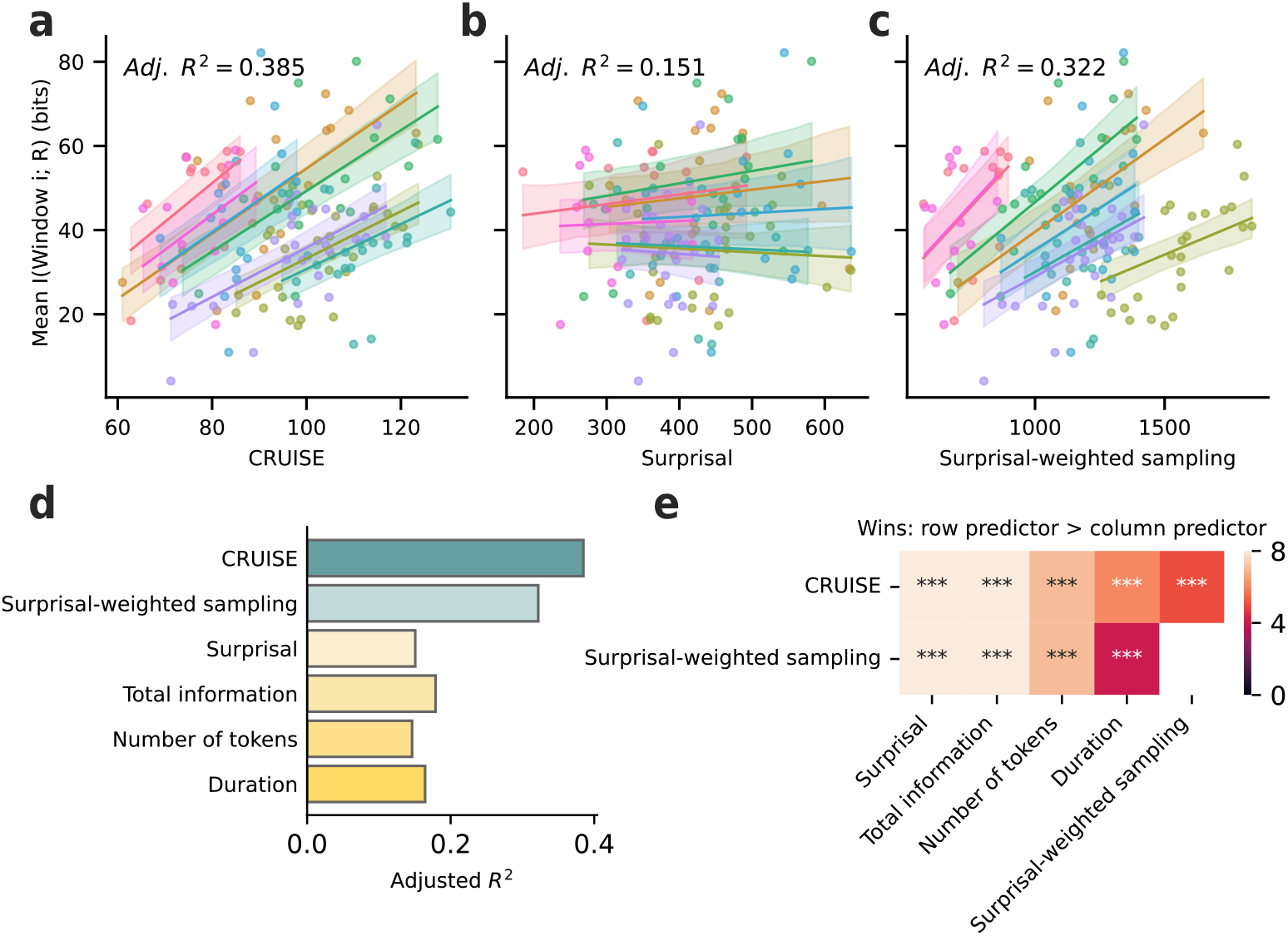
Humans uniformly sample information in time. **a.-c.** Models predicting the mean amount of information participants’ recall captured about each equal-duration text window using **a**. CRUISE, **b**. surprisal, as predicted by event segmentation theory, and **c**. surprisal-weighted sampling. Shaded areas indicate 95% confidence intervals. Each marker is one story window, with different colors for different stories. **d**. *R*^2^ of models predicting the mean amount of information participants’ recall captured about each equal-duration window using linear regression. Each predictor is fitted in a separate linear regression model. Each story is allowed to have its own slope. CRUISE predicts memory the best, followed by surprisal-weighted sampling. **e**. Significance testing of differences between models. Colors indicate the number of stories (out of 8) in which the row predictor predicts significantly more participants’ recalls than the column predictor. Stars indicate the significance of the second-level binomial test: whether the number of stories in which the row predictor significantly better predicted more participants than the column predictor is greater than chance. CRUISE significantly outperforms all other models.

We also compared CRUISE and surprisal-weighted sampling to several controls. It is possible that people remember more about windows in the story that contain more information, or are simply longer. To test these possibilities, we calculated the amount of total information in each window (*H*(*X*_*i*_)), the length of each window in tokens, and the length of each window in seconds. We then fit linear regression models separately for each of these predictors, allowing each story to have a different slope as before. None of the control models’ fit exceeds the fit of either CRUISE or surprisal-weighted sampling (Fig. 2d, total information: *Adj*.*R*^2^ = 0.179, duration: *Adj*.*R*^2^ = 0.164, number of tokens: *Adj*.*R*^2^ = 0.146, also see Fig. S5, and results from other LLMs in Fig. S4).

To statistically compare these models, we performed a slightly different analysis. First, we tested how well each model predicted each individual participant’s recall by computing the correlation between the predictor and participant recall values *I*(*R*^*Y*^, *X*_*i*_). A one-tailed binomial test was then used for each story and pair of models to assess whether one model consistently yielded higher correlation than the other across participants. Finally, we used a second one-tailed binomial test across stories to assess whether the number of stories in which one model outperforms another was greater than chance level. This procedure was used to compare CRUISE and surprisal-weighted sampling to surprisal and control models, and to compare CRUISE to surprisal-weighted sampling. Results show that both CRUISE and surprisal-weighted sampling significantly outperformed surprisal and the other controls (Fig. 2e). CRUISE also significantly outperformed surprisal-weighted sampling.

Overall, these results support our hypothesis that participants sample uniformly from natural language. We replicated these analyses using consensus events segmented by participants (Fig. S7, Methods: Behavioral data analysis), showing that our evidence for the uniform incremental sampling strategy is robust to how the continuous narrative is divided.

### 2.4 Uniform incremental sampling predicts memory at event boundaries

Given that CRUISE outperforms the surprisal model, we next tested whether CRUISE explains memory at event boundaries (wherein one “coherent situation” ends and the next one begins), which is what event segmentation theory was developed to explain. Past research has found that information around event boundaries is better remembered than information within an event [31, 32, 33, 6]. Event segmentation theory proposes that event boundaries are better remembered because surprise at boundaries triggers extensive processing to prioritize encoding [5]. This process requires both feedforward processing, in which one uses long-term memory to compare expectation with the incoming information, and a feedback mechanism to prioritize the encoding of surprising information. We propose that this explanation is unnecessary, as non-uniform memory at boundaries can arise from uniform sampling if boundaries contain more shared information, a feedforward-only mechanism. If true, CRUISE should predict higher memory at event boundaries and knowledge about when boundaries occur should explain no additional variance in behavior.

To test whether CRUISE explains superior memory around event boundaries, we first divided stories into windows of equal length in tokens (Fig. 3a). Windows containing one or more event boundaries were designated “boundary” windows, and those without as “inner” windows. To balance the number of boundary and inner windows while minimizing the number of windows containing multiple boundaries, we chose the number of windows to be 1.5 times the number of events in each story. The edges of each window were minimally adjusted to match phrase boundaries (See Methods: Memory at event boundaries). Note that here we divided the story into windows of even number of tokens instead of duration as before because windows of even number of tokens have approximately equal information. Results using windows of equal duration (Fig. S10, Supplemental results: Alternative event boundary analysis) show a consistent pattern with these results.

**Figure 3.**
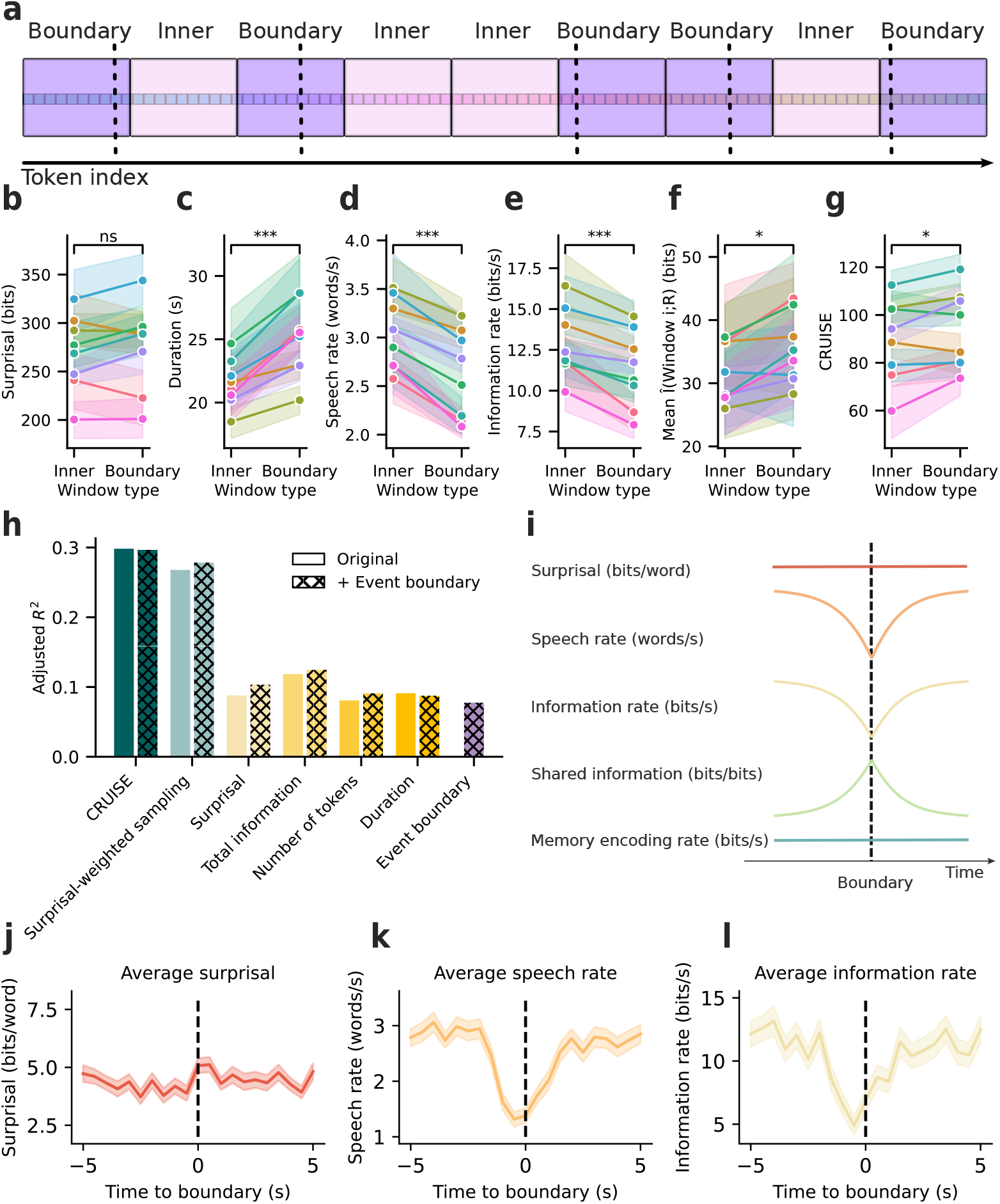
Uniform incremental sampling predicts memory at event boundaries. **a.** We divided stories into windows of equal length in tokens, minimally adjusted to phrase boundaries. The number of windows is 1.5 times the number of events. “Boundary windows” contain at least one event boundary; “inner” windows do not. **b**. Boundary and inner windows do not differ in surprisal (*p >* 0.2, shading indicates 95% confidence intervals. Colors correspond to different stories.) **c**. Boundary windows have significantly longer duration (*p* = 5.85 × 10^−10^), **d**. significantly lower speech rate (*p* = 5.14 × 10^−9^), **e**. significantly lower information rate (*p* = 2.83 × 10^−5^), **f**. and are significantly better recalled than inner windows (*p* = 0.043). **g**. CRUISE predicts that boundary windows are significantly better recalled than inner windows (*p* = 0.015). **h**. *R*^2^ of predicting mean recall using linear regression. Crosshatched bars show models with explicit event boundary information. Boundaries do not explain additional information on top of CRUISE. **i**. Schematic of information properties at event boundaries. Surprisal remains constant but speech rate decreases, lowering the information rate. The amount of shared information increases, enabling better memory for boundaries with constant memory encoding rate. **j.-l**. Average surprisal, speech rate, and information rate around event boundaries in 500 ms bins. Shaded areas indicate standard error. Boundary (time 0) occurs at the right edge of the center bin. Mutual information estimates are not meaningful at the word level, so shared information and memory rate 1w0ere not computed. At event boundaries, **j**. Surprisal remains constant, **k**. speech rate decreases, and **l**. information rate decreases.

We next examined the information properties of event boundaries. While boundary windows and inner windows do not differ in the number of tokens or surprisal (*p >* 0.2), countering the prediction of event segmentation theory (Fig. 3b), boundary windows have significantly longer duration than inner windows (Fig. 3c, *F* = 42.153,*p* = 5.85 ×10^−10^). This reflects lower speech rate (Fig. 3d, *F* = 37.117,*p* = 5.14 ×10^−9^) and lower information rate (Fig. 3e, *F* = 18.318,*p* = 2.83 ×10^−5^) in boundary windows, violating the uniform information density commonly observed in language [27, 24, 34]. Furthermore, by averaging the time courses of surprisal (Fig. 3j), speech rate (Fig. 3k) and information rate (Fig. 3l) around event boundaries, we found that rather than an increase in surprisal, there is a sharp decrease in speech rate and information rate within 5 seconds of event boundaries. Comparing the surprisal around event boundaries to randomly sampled windows from stories, we again confirmed that boundaries are not more surprising (Fig. S11). These results argue against the surprisal account of event segmentation theory.

Next, we tested whether event boundaries were better remembered in our data. A two-way ANOVA (Window type × Story) confirmed that the main effect of window type was significant (Fig. 3f, *F* = 4.155,*p* = 0.043), meaning that participants recalled more information in boundary windows than inner windows (mean boundary *I*(*X*_*i*_, *R*) = 34.26 bits, mean inner *I*(*X*_*i*_, *R*) = 31.14 bits). The main effect of story (*F* = 4.06,*p* = 0.0003) was significant, while the window type × story interaction was not (*p* = 0.938). (Also see Supplemental results: Alternative event boundary analysis).

We next tested whether our uniform incremental sampling strategies predicted participants’ recalls for these boundary and inner windows. As before, we fit linear regression models separately for CRUISE, surprisal-weighted sampling, and controls to predict the mean amount of information that recalls captured about each window (Fig. 3h). Each story was allowed to have a separate slope. Results show that CRUISE and surprisal-weighted sampling explain more variance in *I*(*X*_*i*_, *R*) than surprisal and other control models, with CRUISE having the best performance (CRUISE: *Adj*.*R*^2^ = 0.298, surprisal-weighted sampling: *Adj*.*R*^2^ = 0.268, surprisal and controls: *Adj*.*R*^2^ *<* 0.12). We next asked how CRUISE and surprisal-weighted sampling were able to account for participants’ memory at boundaries. CRUISE could predict stronger memory at boundaries in two ways: first, if event boundaries have higher shared information than non-boundaries; and second, by storing more bits at boundaries due to the lower speech rate and longer duration. Surprisal-weighted sampling, on the other hand, can only predict the stronger memory at boundaries if shared information is higher, because surprisal does not differ between boundaries and non-boundaries. The fact that both CRUISE and surprisal-weighted sampling predicted participants’ recall suggested that boundary windows have higher shared information. The contribution of shared information at boundaries is further corroborated by duration poorly predicting recall, even though boundary windows are significantly longer.

Next, we conducted a two-way (Window type × Story) ANOVA to test whether CRUISE and surprisal-weighted sampling predict that participants better remember boundary windows than inner windows. Only CRUISE predicts that boundary windows are significantly better remembered than inner windows (Fig. 3g, significant main effect of window type, *F* = 5.993,*p* = 1.51 × 10^−2^, non-significant window type × story interaction, *p>* 0.1), while surprisal-weighted sampling does not (main effect of window type *p >* 0.3). These results suggest that shared information alone is not sufficient to predict memory at event boundaries: explicitly considering the slower speech rate and information rate at event boundaries is also crucial.

It is possible that even though CRUISE predicts memory at event boundaries, there is still additional variance captured by the presence of boundaries that is not explained by CRUISE. To test this possibility explicitly, we added either an intercept term for window type (Inner vs. Boundary), an interaction between the continuous predictor and window type, or both (Fig. 3h). A significant intercept term would suggest that participants’ memory still differs for boundary versus inner windows even after accounting for CRUISE. A significant interaction between window type and CRUISE would suggest that the rate of sampling differs for boundary and inner windows. However, we found that neither the intercept nor the interaction explains additional variance on top of CRUISE (Window type intercept only: *p* = 0.25, interaction of window type with CRUISE only: *p* = 0.295). The window type intercept marginally explains additional variance on top of surprisal-weighted sampling (intercept only: *p* = 0.051, interaction only: *p* = 0.113). When both intercept and interaction are added to the regression models, neither is significant for CRUISE or surprisal-weighted sampling (*p >* 0.1). These results suggest that CRUISE fully accounts for any variance in participants’ memory that is explained by event boundaries, but surprisal-weighted sampling does not.

So far, we know that event boundaries are better remembered because they have lower information rate and higher shared information, but does the meaning of information change at event boundaries? Here we used mutual information to quantify the changes in semantic meaning between pairs of windows within an event versus pairs on different sides of an event boundary (Fig. 4a, see Methods: Mutual information within event vs. across an event boundary). We found that on average, within-event pairs have higher mutual information than pairs across a boundary (Fig. 4b, mean per-token mutual information between an across-boundary pair = 0.100 bits, between a within-event pair = 0.324 bits, a 3-way story × distance bin × pair type (across one boundary vs. within-event) ANOVA showed a significant main effect of pair type, *F* = 12.652,*p* = 3.79 × 10^−4^), even though the distances between segment pairs were the same (see Methods: Mutual information within event vs. across an event boundary). This difference in mutual information within an event versus across a boundary was robust to the number of distance bins (six, seven, and eight), and to a normalized shared information measure that corrects for the mean cross entropy of each specific token. Across all variations, 3-way ANOVAs showed higher per-token shared information within-event than across a boundary (*p <* 0.001). These results suggest that events are coherent in their semantic content, and event boundaries indicate shifts in such coherence.

**Figure 4.**
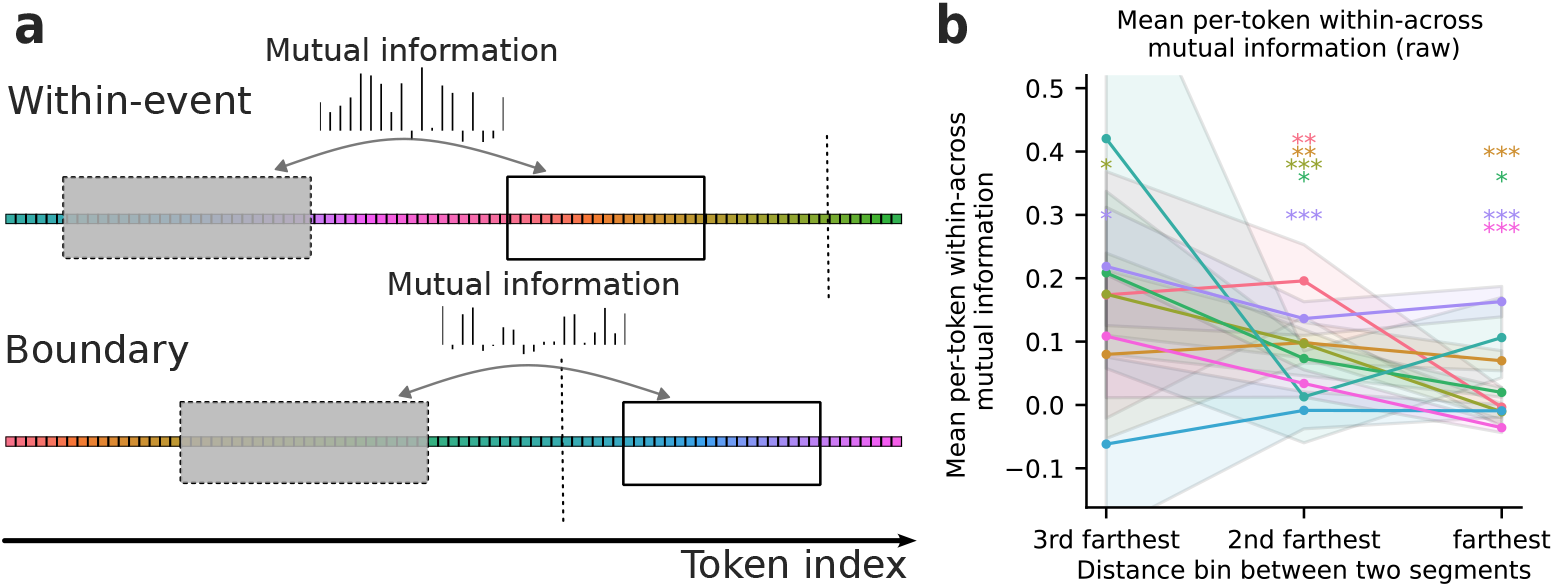
Events are coherent in their semantic content. **a.** Mutual information is calculated for segment pairs within an event versus across one event boundary. A segment pair belongs to the within-event condition when there is no event boundary from the start of the first segment to the end of the second segment. A segment pair belongs to the boundary condition if there’s exactly one event boundary between the end of the first segment and the start of the second segment, and there are no boundaries within the second segment. **b**. The difference between the mean amount of mutual information for within-event pairs and boundary event pairs, separated for stories and distance between two segments. Considering the eight stories together, within-event pairs had higher mutual information than boundary pairs (*F* = 12.652,*p* = 3.79 × 10^−4^). Stars indicate the FDR-corrected significance of post-hoc Welch’s t-tests comparing whether mutual information differs for within-event and boundary pairs for each story and each distance bin.

Overall, these results suggest that stronger memory at event boundaries can be explained by uniformly sampling the incoming information at a constant rate (Fig. 3i, teal line). Thus, event boundaries are better remembered not because they are more surprising (Fig. 3b, j) or sampled at a higher rate by the listener, but because they share more information with the rest of the story as implied by CRUISE (Fig. 3i), and are delivered at a slower speech rate (Fig. 3k), yielding a lower information rate (Fig. 3l). The increase in shared information and decrease in information rate at boundaries are accompanied by a shift in semantic meaning. These results argue against the surprisal account of event segmentation theory.

### 2.5 How do humans form a coherent memory of the narrative?

The preceding results suggest that by utilizing shared information, uniform sampling explains why some parts of a narrative are better remembered. But how do we form a coherent memory of the entire narrative? Selective encoding strategies like event segmentation theory do not offer an explanation because they only focus on how discrete parts of a narrative are remembered. One way to conceptualize how a coherent memory is formed is to view memory as a lossy compression problem, in which the source (a narrative) is compressed into memory imperfectly. The standard framework for this type of problem is rate-distortion theory [35], which was developed to understand lossy compression in communication. Given limited capacity of the information channel (in this case, human memory) and noise in the transmission process, one cannot spend infinite number of bits and distortion is inevitable. Rate-distortion theory suggests that, for a given source and a fixed distortion *d*, which measures how much the compressed memory deviates from the story, there exists a minimal achievable rate *r*(*d*), the number of resources in bits that are used to remember the stimulus, representing the best-case scenario. The function *r*(*d*) is represented as a rate-distortion curve (Fig. 5a, black line). *r*(*d*) is a monotonically decreasing function of *d*, suggesting a rate-distortion trade-off: one can only achieve lower distortion by spending higher rate. Systems of different efficiencies are manifested as different rate-distortion curves: more efficient systems (blue curve) could achieve the same distortion with a lower rate, while less efficient systems (red curve) need a higher rate.

**Figure 5.**
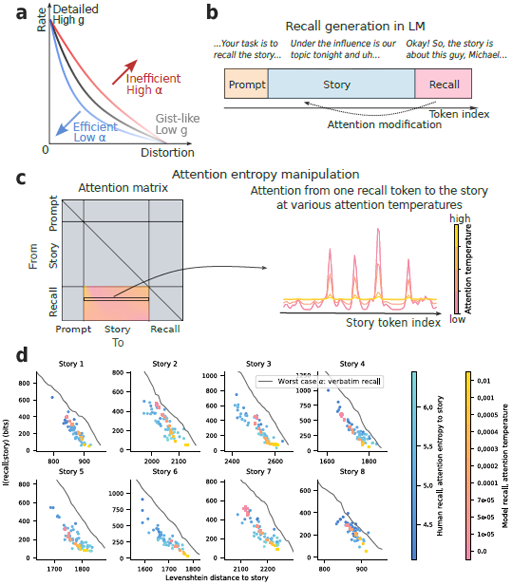
Participants vary along gist vs. detail at similar levels of efficiency. **a.** Based on rate-distortion theory, rate is the number of resources one spent on memorizing the stimuli. Distortion is the deviation between one’s memory and the stimuli. With less efficient memory (larger *α* in our model), one needs larger rate to achieve the same level of distortion. At each level of efficiency, varying the sampling rate *g* trades off rate with distortion. **b**. We prompted an LLM to generate recalls of the same stories heard by our participants. **c**. To manipulate *g* in LLMs, we modified the attention weights from each token in the recall to the story using additive smoothing, while leaving other attention weights intact. The degree of smoothing is controlled by an “attention temperature” parameter, where a value of 0 indicates no smoothing, and higher temperatures make the attention more uniform. **d**. Rate-distortion plots of individual human recalls (blue) and model-generated recalls averaged within each attention temperature (orange-pink). Error bars for model-generated recalls indicate the standard error of all recalls for an attention temperature. We operationalized rate as the mutual information between the recall and the story, and distortion as Levenshtein distance. Human recalls are colored by the mean attention entropy of an induction head from the recall to the story, indicating the level of detail (darker) vs. gist (lighter). Model-generated recalls are colored by the attention temperature used for generation. For both humans and the LLM, more detailed recalls have lower distortion but higher rate. Gray line represents simulated recalls of an individual with no knowledge of the English language, corresponding to the worst-case *α* in CRUISE.

To apply rate-distortion theory to our data, we operationalized rate as the mutual information between story and recall, *I*(*X*; *R*), and distortion as the Levenshtein distance between story and recall, which measures the minimum number of word insertions, deletions, or substitutions required to transform the recall transcript into the story. As expected, the human recalls show a rate-distortion tradeoff (Fig. 5d). However, rate-distortion theory does not explain what information is being lost as distortion increases. Uniform sampling theory, on the other hand, predicts that shared information is effectively oversampled (Eq. 5). Thus, as rate decreases, information shared across the entire story (gist) will be automatically preserved while unique details are lost. If participants adopted the uniform sampling strategy, then recalls with low rate and high distortion would seem more gist-like, summarizing the story. Otherwise, at a low rate and high distortion, participants could recall parts of the story in high detail but have no memory of other parts. In line with predictions of uniform sampling, recalls with varying levels of detail versus gist fall along a rate-distortion curve (Fig. S13). Recalls with a lower rate and a higher Levenshtein distance were rated by human annotators as more gist-like (Fig. S14a,b, *ps <* 0.001). These results suggest that participants adopted uniform sampling, which efficiently compresses the story and automatically extracts the gist, instead of selectively remembering few parts of the story in high detail. The variation in gist versus detail in one’s recall is then achieved by trading off rate and distortion.

Given that uniform sampling explains what information is remembered as we trade off rate and distortion, how does uniform sampling determine where one lies on the rate-distortion plot? To answer this question, we re-expressed the uniform sampling model to explain how the entire story is remembered. Under uniform sampling, the amount of information one remembers about a given story, or rate, *I*(*X*; *R*), is simply the total number of samples that fall within *H*(*X*):

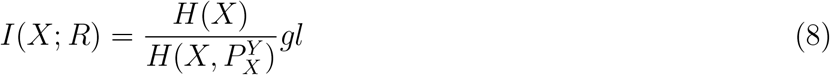

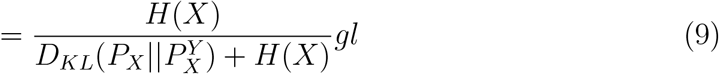

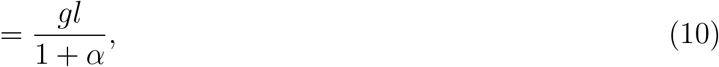

in which *g* is still the constant sampling rate in bits per second, and *l* is the duration of the entire story in seconds. We again assume 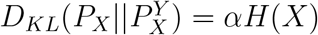, in which *α* is a positive constant. Thus, for a given story *X*, how much one remembers about the story only depends on two factors: the sampling rate *g* and how inadequately one’s prior knowledge captures the stimuli, *α*. We hypothesize that *α* and *g* predict two independent facets of rate-distortion theory: the shape of the rate-distortion curve and where one lies on the curve.

In rate-distortion theory, the shape of the rate-distortion curve is determined by how the information coding scheme maps to the source stimulus. In the case of human memory, this is exactly equivalent to how well the listener’s existing knowledge 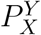 captures the statistics of the stimulus *P*_*X*_, determined by *α* in our uniform sampling model. Once we established the shape of the rate-distortion curve, for a given source and coding scheme, one’s position on the curve is then limited by the channel capacity. One cannot achieve a rate higher than the channel capacity, which is *gl* in the uniform sampling model. Because *l* is fixed for a given story, the sampling rate *g* controls capacity, thereby controlling one’s position on a rate-distortion curve.

### 2.6 Prior knowledge influences memory efficiency

We first validated whether *α* affects the shape of the rate-distortion curve, equivalent to one’s memory efficiency. We did not directly measure or manipulate *α* in our participants, but we simulated the worst-case scenario of *α* to demonstrate how prior knowledge affects the shape of the rate-distortion curve. The worst-case *α* represents a person with no knowledge of the English language trying to memorize the stories. This person could not exploit shared information to compress information, so they would have to memorize the entire story word by word. We mimicked this condition by creating “verbatim recalls” that were simply samples from the story of varying lengths.

We compared the memory efficiency under this worst-case *α* with the memory efficiency of our participants. A memory is more efficient if it uses a smaller rate to achieve the same level of distortion, or, equivalently, achieves lower distortion with the same rate (Fig. 5a). Thus, if prior knowledge determines memory efficiency and participants exploited prior knowledge (smaller *α*) to remember the stories, they would lie below the rate-distortion curve of the worst-case *α*. In Fig. 5d, the rate-distortion curve for the worst-case *α* (gray line) is indeed above recalls for all stories, except for 3 human recalls in Story 8. This result supports our hypothesis that *α*, which measures how different one’s prior knowledge is from the story, affects the shape of the rate-distortion curve. Moreover, it shows that our participants used prior knowledge to efficiently compress the stories into memory.

### 2.7 Rate of sampling predicts the variation of gist versus detail

We next tested whether *g* controls the rate-distortion tradeoff as predicted by our uniform sampling model. While we did not measure the sampling rate behaviorally, we could model the sampling rate using LLMs. When predicting a new token, LLMs take weighted combinations of information from previous tokens where the weights are computed by dot-product attention[36]. The entropy of these attention weights is thus related to how much information can flow from past inputs (the story) to future predictions (recall of the story) [37]. At low attention entropy, each recall token can draw from very specific story tokens, effectively sampling from the story at a high rate. Conversely, at the highest attention entropy, information is simply averaged across the story tokens, effectively sampling at a very low rate. Attention entropy should thus be inversely proportional to the sampling rate *g* in humans, both determining channel capacity.

If the sampling rate *g* controls the rate-distortion tradeoff and the level of detail in one’s recall, so should our proxy of *g*, the attention entropy. To test this hypothesis, we focused on a particular type of attention head inside the LLMs called induction heads [38], which were found independently (see Methods: Evaluation of the attention entropy of induction heads). We concatenated the story and the recall, and calculated the attention entropy from each recall token to the story, then averaged the attention entropy across all recall tokens. We found that recalls corresponding to high attention entropy in the strongest induction head are rated to be more gist-like, having low rate and high distortion, and vice versa (Fig. S15,S14c). These results suggest that sampling rate correlates with the level of detail in one’s recall and the rate-distortion tradeoff. This also shows that it may be possible to quantify how gist-like a person’s recall is by measuring the attention entropy.

Finally, to further test whether the sampling rate *g* causally drives the rate-distortion tradeoff, we generated recalls while manipulating *g* in LLMs, but keeping *α* constant. If *g* drives the rate-distortion tradeoff as predicted by the uniform sampling model, increasing the attention entropy (equivalent to decreasing *g*) should make the LLM generate more gist-like recalls with low rate and high distortion. To do this, we prompted an LLM to generate recalls after “reading” the story (Fig. 5b). We manipulated the attention entropy while preserving the ability of LLMs to generate coherent recalls by applying smoothing to the attention weights from recall tokens to story tokens (Fig. 5d). This process contains one parameter, the “attention temperature” (see Methods: Attention entropy manipulation). An attention temperature of 0 represents the original model without smoothing (low attention entropy), in which the attention heads can precisely sample information from each story token, akin to a high sampling rate *g*. As we increase the temperature, attention entropy increases (Fig. 5c), reducing LLMs’ ability to sample specific information, akin to a low *g*.

Indeed, at temperature 0, the generated recalls of the Llama3-8b-instruct model tended to be rather detailed, often quoting exact expressions from the story (Fig. 5d, also see example generations in Supplemental Tables S2,S3). As we increased the attention temperature, akin to lowering *g* and restricting the channel capacity, the generated recalls used fewer bits at the cost of higher distortion (Fig. 5d), similar to gist-like human recalls. These results suggest that manipulating the sampling rate *g* drives the rate-distortion tradeoff. Further, generating recalls at a high attention temperature explicitly enforces uniform sampling with a low sampling rate, again demonstrating that uniform sampling automatically extracts the gist of a story. Moreover, manipulating attention temperature in LLMs not only drives the rate-distortion tradeoff but also generates human-like recalls in terms of their rate. In seven out of eight stories, we found attention temperatures in which the model-generated recalls matched the human recalls using the same generation prompt. For the remaining story, we also found matching attention temperatures when prompting the LLM to generate more detailed recalls (Fig. S16). These results suggest that LLMs are capable of producing human-like recalls.

Overall, these findings suggest that uniform sampling not only explains why parts of the story are better remembered, but also how a coherent memory of a narrative is formed. By sampling uniformly, participants automatically extract the gist of the story by effectively oversampling information shared across the story. Participants’ memory efficiency, or the shape of their rate-distortion curve, is influenced by how inadequately one’s prior knowledge captures the stimuli, *α*. For a given *α* and a given story, the level of detail versus gist in participants’ recalls, or the position along a rate-distortion curve, is determined by the sampling rate *g*, which controls the capacity of the memory channel. While we did not measure *α* and *g* behaviorally, future work should manipulate participants’ prior knowledge and their sampling rate to test their contribution to memory.

## 3 Discussion

In this study we found evidence that humans uniformly sample information from continuous naturalistic stimuli into memory. While existing theories only address why some parts of a narrative are better remembered, uniform sampling can explain both non-uniform and gist memory by effectively oversampling shared information. Our theory also predicts that a person’s ability to utilize shared information depends on how well their prior knowledge captures the stimulus statistics. The level of gist versus detail is then determined by one’s uniform sampling rate. Overall, our uniform sampling model is mechanistically parsimonious, eliminating the need for a feedback mechanism to decide whether to encode or discard information. These results replicate across multiple narrative stories with distinct themes, and are robust against various methods for quantifying stimulus structure. Our approach extends information-theoretic perspectives that have been used to show that humans efficiently perceive [14, 15, 16, 39] and remember [17, 18, 19, 20, 21] by compressing redundant information. This information-theoretic model uniquely accounts for how non-uniform memory and gist extraction arise from a single computational mechanism.

Our theory is a simpler account of human memory than even segmentation theory, which holds that humans better remember event boundaries due to a feedforward-feedback mechanism that detects surprisal and resets event models, triggering prioritized encoding [5, 32]. That theory arose from a series of experiments using manipulations wherein stimulus change is used as a proxy for surprise [6, 40, 4, 41], or interrupted continuous viewing to ask for subjective surprisal judgments, invoking task demands that may have altered perception of surprise [32] (c.f. [42, 43]). However, experiments using natural narratives have found that surprisal as measured by language models poorly predicts the perception of event boundaries [7]. Our results show that event boundaries are, in fact, not more surprising than non-boundaries (Fig. 3i, red line). Rather, boundaries are simply shifts in semantic content, such that mutual information is higher within an event than across an event boundary. This aligns with earlier results connecting event boundaries to semantics [3, 44, 45]. Our results also align with recent work showing better memory for events with stronger causal and semantic centrality [22, 23]. However, those experiments used human annotations to identify causal connections and text embedding distances to quantify semantic connections. Our model simplifies these computations by directly measuring mutual information using large language models, capturing both semantic and causal connections in a single measure. Mechanistically, event segmentation theory suggests that event boundaries elicit a surprise signal through feedforward comparison, which feeds back to hippocampus to prioritize encoding [46, 32]. In contrast, our theory proposes a parsimonious, feedforward-only mechanism, wherein uniform sampling alone can account for why some moments, including event boundaries, are better remembered. We suggest that event boundaries share more information with the rest of the story than non-boundaries (Fig. 3i, green line). Thus, information at event boundaries is effectively oversampled even with uniform sampling in time. We also found that speech rate is slower at event boundaries, leading to a lower information rate (Fig. 3i, orange and yellow lines). Slower speech rate makes more samples available for encoding at event boundaries than non-boundaries, despite the constant sampling rate (Fig. 3i, teal line) and surprisal (Fig. 3i, red line).

Our theory also provides a straightforward account of the experimental observation that less recent boundaries are better remembered than more recent ones [6]. Because surprise alone cannot account for these temporal differences in memory, event segmentation theory proposes an additional, computationally expensive process wherein working memory is reset at boundaries. In contrast, uniform sampling explains this effect by proposing that memories of earlier boundaries are strengthened as one experiences other parts of the narrative that share information with those earlier boundaries.

Importantly, because uniform sampling effectively oversamples information that is shared across a narrative, it also automatically extracts the gist of a story, a process not accounted for by event segmentation theory. Variation in the level of detail versus gist in one’s recall is then controlled by the sampling rate, *g*. While the level of gist was traditionally measured using subjective human ratings, we also present a novel, theory-driven measure of gist: the attention entropy of induction heads in an LLM. Attention entropy quantifies how specifically each part of the recall corresponds to the narrative. Low attention entropy reflects a verbatim recall of the narrative, while high attention entropy corresponds to gist-like recall. Our method provides an automatic and easy-to-compute measure of gist, opening up possibilities for future work that could rigorously examine neural mechanisms of gist recall without expensive human ratings. Further, we showed that LLMs could achieve human-like recall by manipulating the attention temperature. These results add to previous literature showing that LLMs can produce human-like event segmentation patterns [47, 48]. Besides the sampling rate *g*, our theory also predicts that one’s memory efficiency depends on *α*, or how well one’s prior knowledge captures the statistics of the stimuli. Future studies could directly study the effects of *α* and *g* by comparing participants with varying levels of prior knowledge (e.g. adult versus children, experts versus non-experts), or instructing participants to produce more detailed or gist-like recalls. Our theory also offers a novel quantitative framework for exploring how prior knowledge *α* influences the sampling rate *g* through hippocampal-cortical crosstalk. Overall, our theory provides a major advancement in narrative understanding and opens several new directions for empirical research.

Uniform sampling theory also has implications for communication as an interactive process. Linguistic theories argue that speakers make optimal use of limited communication bandwidth [10, 9] by modulating their speech rate and word choice to achieve approximately uniform information density over time [24, 25, 26, 27, 28, 49]. However, information density is known to be locally non-uniform [26, 50]. We argue that local non-uniformity could be explained if speakers are actually optimizing for efficiency of the listener’s long-term memory system. This would suggest that speakers slow down not only during moments of high information, but also during moments of high shared information, making more samples available for encoding crucial content. Our results support this, showing that event boundaries have lower speech rate and information density but higher shared information, allowing listeners’ memory sampling rate to remain constant. Existing literature in sentence processing has long suggested that processing cost scales with listeners’ working memory demand [49, 8, 51, 52]. Here we extend those ideas to the level of narratives. Future research should test whether modifying the shared information structure and the information rate using LLMs could manipulate what one remembers in a targeted manner.

Our work highlights the importance of considering the information structure of the stimuli [53], thereby accounting for natural statistics of the stimulus modality and the intention of the speaker. One important question for future research is whether our theory extends to other stimulus modalities. For instance, language and visual stimuli have different statistical properties that could interact with memory. Commercial movies are intentionally edited to orchestrate interest and agreement among viewers [54], which could lead to non-uniform information density. While our uniform sampling theory could recover the stimulus structure efficiently for language stimuli with locally non-uniform information density, it is possible that humans could employ alternative strategies to selectively encode information in other modalities. Future work should explore whether humans uniformly sample information in visual or multi-modal stimuli. As the field of artificial intelligence continues to develop multi-modal models [55, 56, 57], we can better capture the statistics of multi-modal stimuli. One limitation of the current study is that our estimates of entropy and mutual information are not perfect. Although LLMs are the state-of-the-art at capturing language statistics, they nonetheless provide only an upper-bound estimate of cross entropy. LLMs achieve superhuman performance in next-word prediction [58, 59, 60], suggesting the upper-bound is lower than humans’ estimates. Yet here they allow us to quantify the interaction between stimuli structure and human memory, a previously intractable problem. To ensure we obtained the best possible mutual information estimates, we selected the method of estimation based on two theoretical properties of mutual information: symmetry and non-negativity. We also replicated our results across five LLMs, and showed that prior exposure to the stimuli did not affect our results by fine-tuning Llama3.2-3b-inst on 301 stories from *the Moth Radio Hour* (See Methods: Fine-tuning on Llama3.2-3b-inst). With future advances in LLMs, we expect to obtain increasingly accurate information estimates.

Overall, while prior theories of narrative memory only explain memory of discrete moments with a process-intensive mechanism, our uniform incremental sampling model offers a parsimonious mechanism to explain both why certain moments are better remembered, and how humans form a coherent memory of the entire narrative. Our theory substantially extends the current literature on narrative understanding, suggesting that future narrative studies should consider the information structure of the stimuli, the intent of the communicator, listeners’ prior knowledge, and memory capacity. These advancements would not be possible without LLMs, enabling us to estimate the information structure of narrative language.

## 4 Methods

### 4.1 Participants

A total of 413 participants were recruited from the online platform Prolific (216 female, 184 male, 13 non-binary; mean age 40.16 years, s.d. = 12.22 years, not including one participant who misreported their age as 569 years). All participants were native English speakers. Informed consent was obtained in accordance with experimental procedures approved by the Institutional Review Board at the University of Texas at Austin. Participants were compensated at the rate of around $10.14 per hour. The target sample size of 50 participants per story was determined by simulating how much a sample size recovers the consensus event boundaries of a previously collected public dataset [61]. We bootstrapped the dataset for 1000 iterations. On each iteration, the dataset was randomly split into two halves, in which one half (102 participants) was used to calculate the target consensus segmentation, and *n* participants were sampled from the other half without replacement to calculate the sample consensus segmentation. A sample size of *n* = 50 reaches the point of diminishing returns in the number of exact word-level consensus boundaries recovered (65% on average).

Participants who had technical difficulties with audio recordings, whose audio recordings were inaudible, and those who recalled the story with minimal effort (for example, those who only spoke for two sentences or recalled little to no relevant information about the story) were excluded from both the segmentation and the recall analyses. Additionally, two participants were excluded from the recall analyses because they reported forgetting to press the recording button and thus recalled twice. One additional participant indicated that they attempted to take notes during story listening, and were excluded for the recall analyses. For the event segmentation analyses, participants were excluded if they did not segment or answered fewer than 3 out of 5 comprehension checks correctly, which indicated that they paid little attention to the story. Moreover, for Story 1, the first 13 participants were excluded from the recall analysis due to an experiment programming error, in which the recall task was erroneously placed after the comprehension check, potentially biasing participants’ recall. All participants, except for one in Story 4, one in Story 7, and one in Story 8, had not been previously exposed to the stories. These three participants were included in the analyses because the number of times they segmented is within 2SD of the sample. See a full breakdown of participant exclusion in Fig. S1 and the sample size for each story in Table S1. After applying the exclusion criteria, 399 participants were included in at least one analysis (212 female, 175 male, 12 non-binary; mean age 40.12 years, s.d. = 12.08 years, not including one participant who misreported their age as 569 years).

### 4.2 Stimuli

The audio stimuli consisted of eight spoken autobiographical narratives from *the Moth Radio Hour*, ranging from 7 to 14 minutes long. All audio tracks were normalized to −23 LUFS. For the story “Pie Man”, the transcript and timing was obtained from [61]. The rest of the stories were manually transcribed by one listener, and the Penn Phonetics Lab Forced Aligner (P2FA) [62] was used to automatically align the audio to the transcript [63]. Praat [64] was used to manually check and correct the alignment.

### 4.3 Experimental Procedures

Before entering the study, participants were asked to make sure they had at least 20 minutes of uninterrupted time. The experiment contains three sections, story listening, verbal recall, and comprehension check. During the story-listening phase, participants were instructed to press a button on the screen whenever, in their judgment, one meaningful event ends and another begins. They were informed that at the end of the story, they would be asked to recall the story they just heard and answer five multiple-choice questions. They were also instructed not to take notes.

Before the participants listened to the actual story, they were prompted to adjust their volume based on a story from *the Moth Radio Hour* that was not part of the current stimuli set. They then received a practice trial to make sure they understood the segmentation instructions. The practice trial involved a 30-second story narrated by the experimenter. After the practice trial, the segmentation instructions were re-emphasized, and participants were told that there was no right or wrong way to perform the segmentation.

Upon entering the main task, participants listened to the story in one setting, while performing event segmentation. Immediately after the story ended, participants were instructed to verbally describe the story they just heard in as much detail as they could while being recorded. Participants were not allowed to end their recording before a 2-minute timer was up. In the last section of the experiment, participants answered 5 four-alternative-forced-choice questions to test their understanding of the story. These questions were designed such that they could be easily answered if one paid attention to the entire story. They also indicated whether they had heard the story before participating in the experiment.

### 4.4 Behavioral data analysis

#### Event segmentation

For each participant, the timings of their button presses were extracted, and aligned to the last word before each button press. To extract the consensus event segmentation for each story, we calculated the proportion of participants who segmented after each word. The time course was then smoothed with a Gaussian kernel with a standard deviation of 2.5 words. The smoothed time course was then thresholded at 95% percentile, and peaks above the threshold were identified as the consensus event segmentation of the story. This procedure follows an earlier publication [61].

#### Recall transcription

Recall audios were first automatically transcribed using Whisper-large-v3 [65]. Then the automatically generated transcripts were manually checked against the audio recording and corrected for errors.

#### Recall coding

Recall transcripts of Story 1, 2 and 3 were coded using a coding scheme adapted from the Autobiographical Memory Interview (AMI) coding scheme [66] to validate the LLM-derived information measures. Given the number of participants and the length of the recall transcripts, the remaining stories were not coded. Modifications to the AMI coding scheme were made to consider the fact that our participants were recalling the narrator’s autobiographical story, not their own. As in the AMI coding scheme, a recall transcript was divided into individual units named “details”, each conveying unique meaning. Each detail was then assigned to a category. Specifically, all details pertaining to the content of the story itself could be categorized as event, place, time, perceptual, emotion/thought, and semantic details. We added a category of participant-related details to accommodate participants’ reactions and inference of the story. Participants’ thoughts and emotions during the story listening were categorized as emotion/thought-Participant (e.g. “I was really laughing during that part of the story”). Semantic information inferred by the participants was categorized as semantic-Participant. For instance, if the story only mentioned that the narrator went to college in the Bronx, while the participant recalled “in the Bronx, in New York”, “in New York” is categorized as semantic-Participant because it is inferred by the participant. Lastly, repetitions of previously mentioned details were categorized as “repetitions”. Details that do not reflect recollection, such as meta-cognitive statements (“Let me see if I can remember that”), subjective comments about the story (“I like it it’s a really funny story”), inference about the context of story-telling (“it sounded like a comedian was on the stage”), and messages to the experimenter were categorized as “other”.

All details pertaining to the story itself were considered in subsequent analyses. Details categorized as participant-related, “repetitions”, and “other” are excluded. Each detail was marked either as correct or incorrect. We then traced each detail back to its origin in the story, recording the first consensus-derived event in which it appeared. For each correct detail in the recall, we then compared the detail to how it was originally described in the story, and assigned a rating to indicate its level of richness on a scale of 1 to 3. Following the original AMI coding scheme, 3 indicates that the participant’s recall of the detail fully captures the way this detail was described in the original story; 2 indicates that the recall of the detail captures some but not all specific information about how this detail was described in the original story; 1 indicates that the recall only captures general, nonspecific information about the original detail.

Before coding the actual dataset, all coders were trained on the thirteen Story 1 recall transcripts that were excluded from the recall analysis due to an experiment programming error. Three primary coders each coded all recalls for one of the three stories. A single reliability coder coded 10 recalls (about 20% of participants) for each story. A fifth independent coder divided all three stories into details using the same coding scheme above. The primary and reliability recall coders were blind to the story coding during their coding process. After coding was completed, each recall coder was informed of the number of details of each event in the story. Each coder then reviewed their coding to ensure that the number of details recalled per event did not exceed those in the story. The primary coders agree highly with the reliability coder (Cronbach’s alpha of the number of correct details per event: Story 1: 0.98, Story 2: 0.90, Story 3: 0.90; Cronbach’s alpha of the mean level of richness per event: Story 1: 0.91, Story 2: 0.84, Story 3: 0.85).

#### Creating equal-duration windows of story text

To divide the story into windows of approximately equal duration (which we used to evaluate how much participants recalled about each part of the story in Fig. 2), we first split the story into equal-duration windows, in which the number of windows equal to the number of consensus events for each story. The start and end time of each window were then mapped to the corresponding story text. To avoid abnormally high cross entropy due to syntax violations, the beginning and ends of each window were minimally adjusted to match phrase boundaries. For example, a window that starts with “it all so I just blurted out what happened to me” was adjusted so that “it all” was moved to the previous window, so the previous window now ends with “I couldn’t really appreciate it all”, and the current window now starts with “so I just blurted out what happened to me”.

### 4.5 Information estimates

#### Language models

Transformer-based large language models were used to estimate various information measures. These models were trained on large corpora of text to learn the statistics of language, and represent text as subword tokens. We adopted the following language models in this paper: Llama-3-8B-Instruct [29], Llama-3-8B-Base [29], Mistral-7B-Instructv0.3 [67], Llama-3.2-3B-Instruct [68], and Gemma-2-9b-it [69]. All models are openly available on HuggingFace. Inference was performed on a compute node with 3 NVIDIA A100 40GB cards.

#### Estimating total information of a part of a story *H*(*X*_*i*_) and the total information of a recall *H*(*R*)

We used cross-entropy of the language model to estimate the total information in a part of the story. Each part *X*_*i*_ was tokenized using the pre-trained tokenizer and fed separately into the language model without preceding context. For each token in *X*_*i*_, the language model outputs a cross-entropy loss, indicating how inaccurately it predicted the next token based on preceding inputs. Because we used a base-2 logarithm in the cross-entropy calculation, this also gives the number of bits needed to specify the correct next token under the probability distribution of the language model. The total cross-entropy of the part *H*(*X*_*i*_, *Z*) under the language model *Z* is thus the sum of cross entropy of all tokens in the part. Given that 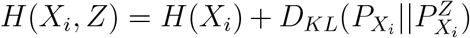 and LLMs are the state-of-the-art at language modeling, we assume that 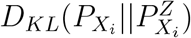 is relatively small and consistent across different texts, and use *H*(*X*_*i*_, *Z*) as an estimate of *H*(*X*_*i*_). Under the same logic, the total information in a recall *H*(*R*) is estimated by the sum of cross entropy of all tokens in the recall.

#### Estimating the surprisal of a part *H*(*X*_*i*_ | *X*_1_,…, *X*_*i*−1_)

The surprisal of a part of the story was estimated by passing the entire story into the LLM, thus presenting each part in context of preceding parts of the story. We then took the sum of cross entropy of all tokens in the part. Because all stories are shorter than the maximum context length of the language models, at each token in a story, the cross entropy output of the language model takes into account the context of prior text in the story.

#### Estimating the mutual information of a pair of parts *I*(*X*_*i*_; *X*_*j*_)

The mutual information of a pair of parts *I*(*X*_*i*_; *X*_*j*_) represents the amount of information reduction in *X*_*j*_ when *X*_*i*_ is provided as context, *I*(*X*_*i*_; *X*_*j*_) = *H*(*X*_*j*_) − *H*(*X*_*j*_ |*X*_*i*_). *H*(*X*_*j*_) is estimated by passing *X*_*j*_ to the LLM and taking the sum of cross entropy of all tokens. *H*(*X*_*j*_ *X*_*i*_) is estimated by directly concatenating the text of *X*_*i*_ and *X*_*j*_, and taking the sum of cross entropy for all tokens in *X*_*j*_ in the context of *X*_*i*_. To avoid potential confusion of the LLMs when parts are presented out of order, which could yield a superfluous 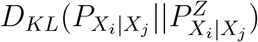, we concatenated the parts according to their order in the story. In other words, for a pair of parts *X*_*i*_ and *X*_*j*_, where *i< j, X*_*j*_ is always concatenated after *X*_*i*_, i.e., *I*(*X*_*i*_; *X*_*j*_) = *I*(*X*_*j*_; *X*_*i*_) = *H*(*X*_*j*_) − *H*(*X*_*j*_ |*X*_*i*_). The mutual information of a part with itself is equivalent to the total information of that part, *I*(*X*_*i*_; *X*_*i*_) = *H*(*X*_*i*_).

#### Estimating the mutual information of recall and a part of the story *I*(*R*; *X*_*i*_)

The mutual information *I*(*R*; *X*_*i*_) of recall *R* and a part *X*_*i*_ represents the amount of information reduction in *X*_*i*_ when *R* is provided as context, or equivalently, the amount of information reduction in *R* when the part *X*_*i*_ is presented in context. Because theoretically mutual information is symmetric, *I*(*R*; *X*_*i*_) = *H*(*X*_*i*_) − *H*(*X*_*i*_ |*R*) = *H*(*R*) *H*(*R* | *X*_*i*_) = *I*(*X*_*i*_; *R*), and there is no explicit ordering of the text of the recall and the part of the story (unlike when there is an order between two parts of the same story), *I*(*X*_*i*_; *R*) can be estimated in both directions: subtracting *H*(*X*_*i*_ | *R*) from *H*(*X*_*i*_), or subtracting *H*(*R* | *X*_*i*_) from *H*(*R*).

To achieve the former, we estimated *H*(*X*_*i*_) by directly passing the part of the story to the model, and taking the sum of cross entropy of all tokens. We then pass the model with a direct concatenation of the recall text with the segment from the story:

~~~
{recall}{segment}
~~~

and took the sum of cross entropy of all tokens in the story segment in the concatenation as *H* (*X*_*i*_ | *R*).

Alternatively, to estimate *I*(*X*_*i*_; *R*) by subtracting *H*(*R* | *X*_*i*_) from *H*(*R*), we estimated *H*(*R*) by directly passing the entire recall to the model, and taking the sum of cross entropy of all tokens. To estimate *H*(*R* |*X*_*i*_), we directly concatenated a segment of the story and then the recall, and took the sum of cross entropy of all recall tokens in the concatenation as *H*(*R* | *X*_*i*_):

~~~
{segment}{recall}
~~~

We directly concatenated the recall and a part of the story in the estimations above. Alternatively, it is also possible that informing LLMs where the text of *R* and *X*_*i*_ begin using prompting can allow LLMs to better estimate the relationship between *R* and *X*_*i*_. To achieve prompting, for instruction-tuned models, we adopted chat templates and split the instructions into system prompts and user prompts. For base models, we simply presented the same instructions without the chat templates. The prompts we used are as follows:

To estimate the total information of a part of the story *H*(*X*_*i*_), we used the following prompt, and summed across the cross entropy of all tokens in *X*_*i*_:

~~~
System prompt: You are going to read a segment from a story.
User prompt: Here’s the segment from the story: {segment text}
~~~

To estimate the total information of a recall text *H*(*R*), we used the following prompt, and summed across the cross entropy of all tokens in *R*:

~~~
System prompt: You are going to read a human’s recall of a story.
User prompt: Here’s the recall: {recall text}
~~~

To estimate the conditional information of the recall text when presented in the context of a part of the story *H*(*R* | *X*_*i*_), we used the following prompt and summed across the cross entropy of all tokens in the recall text:

~~~
System prompt: You are going to read a segment from a story, along
with a human’s recall of the entire story this segment belongs
to.
User prompt: Here’s the segment from the story: {segment text}.
Here’s the recall of the story: {recall text}
~~~

Similarly, to estimate the conditional information of a part in a story given a recall as context *H*(*X*_*i*_| *R*), we used the following prompt and summed across the cross entropy of all tokens in the story segment:

~~~
System prompt: You are going to read a human’s recall of a story, and segment from a story.
User prompt: Here’s the recall of the story:
{recall text}. Here’s the segment from the story: {segment text}
~~~

Thus we have four possible ways to estimate *I*(*X*_*i*_; *R*) (2 concatenation methods: direct concatenation or instruction prompting ×2 concatenation orders: presenting recall text then a part of the story, hereon “recall first”, or presenting a part of the story then recall text, hereon “recall last”). These methods do not give exactly the same estimates. To select the most valid method of estimation, we developed two metrics based on theoretical properties of mutual information.

The first metric assessed the symmetry of *I*(*X*_*i*_; *R*) estimates by taking the root mean square of *I*(*X*_*i*_; *R*) estimated by presenting recall first versus recall last, for all pairs of story parts *X*_*i*_ and recalls *R*_*j*_. This symmetry metric is formulated as follows:

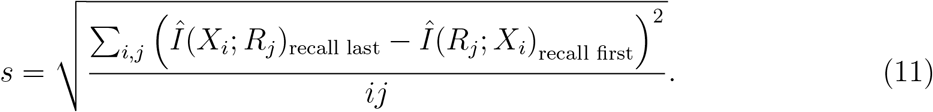

The second metric assessed the theoretical property that mutual information is non-negative. While theoretically *I*(*R, X*_*i*_) is always non-negative, practically, the conditional information *H*(*X*_*i*_|*R*) estimated by a language model could be greater than the estimates of 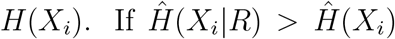, it is a sign that the language model failed to correctly extract the relationship between the recall and a part of the story. The same holds for 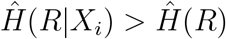. Thus, for each method to estimate *I*(*X*_*i*_; *R*), we calculated the proportion of negative 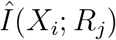 for all pairs of story parts *X*_*i*_ and recalls *R*_*j*_,

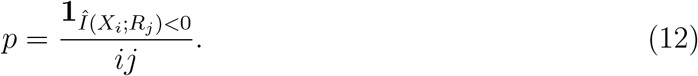

We selected the LLM and estimation method pair based on the symmetry and non-negativity metrics. For both metrics, lower values signal that the mutual information estimates provided by the LLM are more valid. To accomplish our goal of recall generation, a core analysis to investigate the sampling rate *g* (Fig. 5), we needed an instruction-tuned model. We also tested a non-instruction tuned model (Llama3-8b) in Supplemental results (Fig. S4, S8). Across the eight stories, we computed an average of each metric for parts of stories derived from the consensus event segmentation and by splitting the story into windows of equal duration. We first selected the instruction-tuned language model and the concatenation method pair (direct concatenation or prompting) that yielded the lowest root mean square *s* (highest symmetry): Llama3-8b-inst with direct concatenation. We then selected the concatenation order (recall first or recall last) that yielded the lower proportion of negative mutual information estimates. Based on these metrics, we selected Llama3-8b-inst while directly concatenating the recall before the part of the story, and reported those results in the main section of the paper. It might seem odd that an instruction-tuned model provides the best mutual information estimate without instruction prompting. We note that an instruction-tuned model could still preserve the language modeling ability even without instruction prompting, and it is possible that instruction prompting systematically alters the probability distribution output by the LLM, leading to worse mutual information estimates, which is not what LLMs are trained for. We noted that while estimating *I*(*X*_*i*_; *R*) using Llama3-8b base model with prompting and presenting recall first yielded the best *p* and *s* metrics overall (Fig. S2), the base model failed to generate recalls of the story, an essential analysis to uncover the role of sampling rate in Fig. 5. Thus for the coherence of the main results section, results of Llama3-8b base model and others were reported in the Supplemental Results (Fig. S4, S8). All models with reasonable symmetry and non-negativity metrics suggest that CRUISE predicts *I*(*X*_*i*_; *R*) while surprisal and other controls perform poorly.

#### Estimating the mutual information of recall and the entire story *I*(*R*; *X*)

Given that directly concatenating the recall before the part of the story using Llama3-8b-inst yields the most valid mutual information estimates between a part in a story and a recall text, we adopted the same method and order of estimation to estimate *I*(*R*; *X*). In other words, we calculated the change in cross entropy of the story tokens when recall is concatenated before the story compared to when the story is presented to the model alone. This represents the amount of information reduction in *X* when *R* is provided as context, or the amount of information *R* explains about the story *X*.

### 4.6 Linear modeling

We fitted linear regressions to test how well CRUISE, surprisal-weighted sampling, surprisal, and other controls models predicted participants’ memory of each part of the story. For each story, we calculated the mean *I*(*X*_*i*_; *R*) averaged across all participants’ recalls. For each model, we separately fitted a linear regression using data from all eight stories, where the mean *I*(*X*_*i*_; *R*) is the response variable. Each regression has the following predictors: an intercept term, the model of interest, and an interaction term between the model and a categorical variable indicating which story the data point belongs to (i.e. a separate slope for each story). The interaction term was included because participants tended to produce recalls of similar durations regardless of the duration of the story, leading to lower apparent sampling rates for longer stories. Given that the study was conducted online, we believe this reflects a reward-maximizing strategy, instead of a difference in participants’ actual sampling rate and understanding of the story.

### 4.7 Significance testing of CRUISE, surprisal-weighted sampling and controls

We tested whether CRUISE and surprisal-weighted sampling outperformed controls, and whether CRUISE outperformed surprisal-weighted sampling in a pairwise manner. We examined for each pair, whether one model significantly better predicted more participants’ recall than the other. One potential approach is to collapse across participants from all eight stories and perform a one-tailed binomial test with a success probability of 0.5. This approach would yield highly significant results after FDR correction, suggesting that when the stories were split into windows of even duration, both CRUISE and surprisal-weighted sampling outperformed surprisal and other controls, and CRUISE outperformed Surprisal-weighted sampling. When stories were split into consensus events, CRUISE did not outperform surprisal-weighted sampling, but both CRUISE and surprisal-weighted sampling still outperformed surprisal and other controls. However, by observing the data, it is clear that there is between-story variance: recalls of some stories were better predicted by certain models than others. To capture the between-story variance, we conducted a two-level binomial test for each pair of models. At the first level, we tested whether one model significantly better predicted more participants’ recalls than another within each story using a one-tailed binomial test with a success probability of 0.5, and *α* = 0.05. Then, for each pairwise comparison, we counted the number of stories in which the first-level comparison was significant. Then a second-level binomial test was performed to test whether out of the eight stories, obtaining *k* significant stories at the first level was not due to random chance. In other words, at the second-level, we performed a one-tailed binomial test with *H*_0_ : *p* = 0.05 and *H*_1_ : *p>* 0.05, *n* = 8, and *k* = the number of significant stories for that comparison at the first level. Results using the two-level test replicated those from collapsing across all participants’ recalls across eight stories.

### 4.8 Memory at event boundaries

We tested whether the uniform sampling hypothesis explains participants’ memory at event boundaries. We divided each story into windows of even number of tokens. The beginning and ends of each window were then slightly adjusted to the nearest syntax boundaries, so that the language model would not produce abnormally high cross entropy due to syntax violations. The number of windows of each story equals to 1.5 times the number of events. Each window was classified as a boundary window if it contained at least one event boundary, or an inner window if not. The multiplying factor 1.5 was chosen to balance the number of boundary and inner windows in each story, and to minimize the number of boundary windows with more than one event boundary. We estimated *I*(*X*_*i*_; *R*) using each window. To test whether event boundaries explain additional variance beyond CRUISE, surprisal-weighted sampling, or control predictors, we ran a linear regression to predict the average *I*(*X*_*i*_; *R*) across participants. The model contains a target predictor (CRUISE, surprisal-weighted sampling, or a control predictor), an interaction term with the categorical variable story, and either a dummy variable window type (boundary vs. inner), or an interaction between window type and the target predictor, or both. We additionally performed similar boundary analysis by dividing the story into windows of equal duration, and reported the results in the Supplement Results: Alternative event boundary analysis.

### 4.9 Time courses of information properties of event boundaries

We calculated the speech rate, information rate and surprisal around consensus event boundaries in two ways: either for a fixed time window (−5s to 5s around the boundary), or a fixed number of words (5 words before to 5 words after the boundary). Because our consensus event boundaries were determined at the word level, for the fixed time window version in the main text, we took the midpoint between the offset of the last word in the pre-boundary event and the onset of the first word in the post-boundary event as the event boundary. We then created 500 ms bins, where the right edge of the center bin corresponds to the timing of the consensus event boundary. A word is considered to belong to a bin if the midpoint of that word is within the bin edges. The speech rate is computed as the number of words in the bin divided by bin duration; the information rate is the total cross entropy of all words in the bin divided by bin duration; surprisal is the mean cross entropy of all words in the bin. We averaged time courses of all event boundaries across 8 stories. For the version with a fixed number of words in Fig. S12, event boundary (word index 0) is the last word in the pre-boundary event. The speech rate of each word is 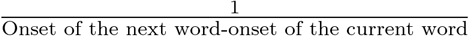; the surprisal of the word is the sum of cross entropy of all tokens in the word; the information rate of each word is 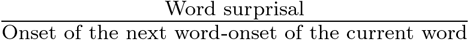.

### 4.10 Mutual information within event vs. across an event boundary

To calculate the mutual information between two segments within an event or across an event boundary, we ablated chunks of text and observed the change in cross entropy of a range of future text at a fixed distance bin from the ablated text. Ablations are generated in a sliding window manner across the entire story. Target lengths of ablations are 10, 30, 50, 70, 90, 110, 130, and 150 tokens. To avoid sub-word ablations yielding superfluous cross entropy, we extended the end of each ablation until the ablation respected word boundaries. Each token was guaranteed to be ablated at least 10 times, except for the final tokens in the story that could not meet the minimum ablation length. For each ablation window, we calculated the amount of mutual information between the ablated window and a window of future text, operationalized as the change in cross entropy of a range of future tokens post-ablation -pre-ablation. A greater difference in cross entropy post-pre ablation indicates that the ablated text has greater mutual information with the range of future tokens. Because cross entropy has a power-law relationship with the ablation distance, we took *k* equally spaced log bins as the ranges of future tokens for which the change in cross entropy was calculated. We separately determined the largest bin edge for each story as the mean consensus event length + 1 standard deviation. We showed results with *k* = 5 log bins in the main text, and replicated the results with *k* = 6, 7, 8. For a pair of ablated text and a range of future tokens to qualify for the within-event condition, there needs to be no event boundary from the end of the ablation to the end of the range of future tokens. For a pair of ablated text and a range of future tokens to qualify for the across one event boundary condition, there needs to be exactly one event boundary between the end of the ablation and the start of the range of future tokens, while there are no event boundaries within the range of future tokens itself. Because semantic relationship emerges over relatively long ranges of text, we only compared the difference of mutual information between within-event vs. across an event boundary conditions for the 3 farthest log bins out of the *k* total bins. Moreover, to correct for the possibility that cross entropy might be differentially affected by ablations due to confounds such as token frequency, we additionally calculated a normalized mutual information measure between the ablated text and the window of future text. To do so, we normalized the change in cross entropy of each token by the mean cross entropy of that token across 301 stories from *the Moth Radio Hour*. Results across all variations replicated those in the main text.

### 4.11 Estimation of rate and distortion

Rate is defined as the number of bits one spent on compressing the stimuli, *I*(*X*; *R*). We estimated *I*(*X*; *R*) by calculating the amount of information reduction in the story given the recall, *I*(*X*; *R*) = *H*(*X*) − *H*(*X* |*R*), see details above. Distortion is operationalized as the word-level Levenshtein distance between the recall transcript and the story transcript. The word-level Levenshtein distance is calculated by mapping each unique word to a unique character, and applying the Python library Polyleven [70]. The Levenshtein distance represents the number of single-word edits required to change the recall text to the story text.

### 4.12 Simulation of the worst case *α*

The worst-case *α* represents the scenario when one has no knowledge of the stimuli, thus they could not utilize the shared information to compress the stimuli, and would have to attempt to remember the story in a verbatim manner. We simulated the rate-distortion curve of this scenario by evaluating the rate and distortion of cumulative recalls, in which recalls incrementally include the first 10, 20, 30 words of the story, and so on.

### 4.13 Evaluation of the attention entropy of induction heads

To model the sampling rate of human recalls, we measured the attention entropy of induction heads from recall tokens to the story. An induction head is defined as an attention head that looks for previous instances of the current token and attends to the token after the previous instance [38]. We independently detected induction heads by concatenating three consecutive presentations of a text segment back-to-back [60]. Thus, in the second presentation, an induction head would look at the token after the previous instance in the first repetition, and in the third presentation, an induction head would look at the corresponding tokens in the first and second repetition. We used the beginnings of stories 1, 2, and 3 as three separate test stimuli. We then calculated, for each test stimulus and each attention head (Llama3-8b-Instruct has 32 layers and 32 heads), the proportion of instances that the head displayed the desired induction head behavior. We then averaged the proportions across three test stimuli to obtain an induction head score. We selected the attention head with the highest induction head score (layer 15, head 30) for further analysis. Code for induction head detection is heavily based on TransformerLens [71].

We then evaluated the attention entropy [37] of the strongest induction head from a recall transcript to the story text. We first concatenated the story and the recall transcript. We normalized the attention from each recall token to all story tokens so that they sum to 1 to control for different amounts of attention to the story versus the recall itself. We next calculated the entropy of the normalized attention distribution from the recall token to all story tokens. We then averaged these values across all recall tokens to obtain the mean attention entropy from recall tokens to story tokens.

### 4.14 Recall generation in language models

We prompted Llama3-8b-inst to “read” the story and generate a recall. We applied the Huggingface chat template with the following prompt:

~~~
System prompt: You are a human with limited memory ability. You’re
going to listen to a story, and your task is to recall the story
and summarize it in your own words in a verbal recording. Respond
as if you’re speaking out loud.
User prompt: Here’s the story: {story text} \nHere’s your recall:
~~~

Because LLM-generated recalls for Story 6 using the above prompt did not reach *I*(*X*; *R*) of human recalls, we subsequently prompted the LLM to generate in more detail for Story 6:

~~~
System prompt: You are a human with limited memory ability. You’re
going to listen to a story, and your task is to recall the story
in your own words in as much detail as you can in a verbal recording.
Respond as if you’re speaking out loud.
User prompt: Here’s the story: {story text} \nHere’s your recall:
~~~

### 4.15 Attention entropy manipulation

To manipulate the degree of gist in the model-generated recalls, we modified the attention entropy from the recall to the story while keeping the attention elsewhere intact. The attention entropy was modified using additive smoothing, while the sum of attention from a recall token to the story remains the same. Let *p*_*ij*_ be the original attention score from recall token *i* to story token *j*. Let 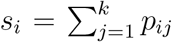 be the total attention from a recall token *i* to all story tokens, in which *k* is the total number of story tokens. Using additive smoothing, we modified the attention from a recall token *i* to a story token *j* to be 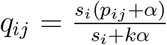. The free parameter *α ∈* ℝ^+^, which we called the *attention temperature*, controls the degree of additive smoothing. When *α* is 0, *q*_*ij*_ = *p*_*ij*_. When 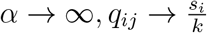, and the attention from the recall token *i* to all story tokens approaches the uniform distribution, thus producing high attention entropy. All attention heads of all layers were manipulated with the same *α*. We then passed the recall generation prompt above to LLMs to generate recalls of the story.

## Supporting information

Supplemental methods and results

## 4.16 Data availability

The data underlying this study will be made publicly available at OSF before publication.

## 4.17 Code availability

The code underlying this study is made publicly available at: https://github.com/HuthLab/CRUISE/tree/main.

## Acknowledgement

We thank Indrani Basu, Trisha Clennan, Karina Kapoor, and Olivia Simmons for assistance with transcribing and coding recall transcripts.

## Notes

### Competing Interest Statement

The authors have declared no competing interest.

