## Supplemental methods and results for "Efficient uniform sampling explains non-uniform memory of narrative stories"

### 5 Supplemental results and figures

#### 5.1 Supplemental Methods

Table S1: Story information and participant exclusion

| | Story name | Duration (min) | Word count | Total participants pre-exclusion | Participants included for segmentation | Participants included for recall |
|---|---|---|---|---|---|---|
| Story 1 | Pie man | 7.5 | 962 | 69 | 58 | 51 |
| Story 2 | Alternate Ithaca Tom | 11.8 | 2174 | 50 | 48 | 47 |
| Story 3 | Ode to stepfather | 13.8 | 2669 | 48 | 42 | 45 |
| Story 4 | Souls | 12.2 | 1867 | 49 | 46 | 47 |
| Story 5 | Legacy | 13.7 | 1891 | 51 | 51 | 49 |
| Story 6 | Where there's smoke | 10.0 | 1836 | 48 | 47 | 46 |
| Story 7 | Adventures in saying yes | 13.4 | 2307 | 49 | 47 | 44 |
| Story 8 | In a moment | 7.2 | 964 | 49 | 48 | 45 |

**Assumptions of CRUISE** In the main text we present a version of CRUISE in which we make a number of simplifying assumptions about how participants' inefficiency is reflected in how inadequately they model each part of the story, $D_{KL}(P_{X_j}||P^Y_{X_j})$. This form accounts for how inadequately participants model the relationship between two parts of the story by diluting the number of samples $X_j$ could contribute to $X_i$ when participants have a greater $D_{KL}(P_{X_j}||P^Y_{X_j})$ (worse model). Making no such assumptions, the full form of CRUISE should be written as

$$I(X_i;R) = \sum_j \frac{H(X_i, P^Y_{X_i}) - H(X_i|X_j, P^Y_{X_i|X_j})}{H(X_j, P^Y_{X_j})} gl_j \quad (13)$$

$$= \sum_j \frac{H(X_i) + D_{KL}(P_{X_i}||P^Y_{X_i}) - H(X_i|X_j) - D_{KL}(P_{X_i|X_j}||P^Y_{X_i|X_j})}{H(X_j) + D_{KL}(P_{X_j}||P^Y_{X_j})} gl_j, \quad (14)$$

in which $P_{X_i}$ and $P_{X_i|X_j}$ are the true distributions of $X_i$ and $X_i|X_j$ respectively, which could be thought of as representing distributions under the model of the speaker or the average listener. $H(X_i, P^Y_{X_i})$ and $H(X_i|X_j, P^Y_{X_i|X_j})$ are the cross-entropy of $X_i$ and $X_i|X_j$ under listener $Y$'s model of the world, respectively. To achieve the simplified form in the main text, we assume that the difference between the KL-divergences in the numerator is negligible, i.e., $D_{KL}(P_{X_i}||P^Y_{X_i}) - D_{KL}(P_{X_i|X_j}||P^Y_{X_i|X_j}) \approx 0$, meaning that participants' models deviate similarly in estimating $H(X_i)$ and $H(X_i|X_j)$. This assumption simplifies the model to:

$$I(X_i; R) = \sum_j \frac{H(X_i) - H(X_i|X_j)}{H(X_j) + D_{KL}(P_{X_j}||P^Y_{X_j})} g l_j \tag{15}$$

$$= \sum_j \frac{I(X_i; X_j)}{H(X_j) + D_{KL}(P_{X_j}||P^Y_{X_j})} g l_j \tag{16}$$

Future work could relax this assumption to explore when participants' models deviate from $X_i$ and $X_i|X_j$ differently, thereby directly capturing how well participants model the relationship between $X_i$ and $X_j$.

We also assume that the cost of representing $X_j$ under listener $Y$'s model is proportional to the amount of information in $X_j$,

$$D_{KL}(P_{X_j}||P^Y_{X_j}) = \alpha H(X_j), \tag{17}$$

in which $\alpha$ is a positive, unitless constant that represents how inefficiently the listener models this story. Intuitively, this reflects the idea that one's difficulty understanding a story will scale with its length or total information. This assumption further simplifies CRUISE to:

$$I(X_i; R) = \frac{g}{1+\alpha} \sum_j \frac{I(X_i; X_j)}{H(X_j)} l_j, \tag{18}$$

allowing us to combine the unknown factors $g$ and $\alpha$ into a single multiplicative constant $\frac{g}{1+\alpha}$ estimated with linear regression. Future work could also consider the scenario in which $D_{KL}(P_{X_j}||P^Y_{X_j})$ is not directly proportional to $H(X_j)$.

**Derivation of surprisal model** In the surprisal model, the total number of bits stored for a part of the story $X_i$ is equivalent to the sampling rate times its surprisal, $gH(X_i|X_1, ...X_{i-1})$. Similar to CRUISE, the number of bits one actually remembers about $X_i$ is also influenced by how inadequately participants model a part of the story. A worse model of a part discounts the number of available samples. Because here participants only sample surprising information within $X_i$, the fraction of useful samples that land within $X_i$ is $\frac{H(X_i|X_1,...X_{i-1})}{H(X_i|X_1,...X_{i-1}) + D_{KL}(P_{X_i|X_1,...X_{i-1}}||P^Y_{X_i|X_1,...X_{i-1}})}$. The number of bits one remembers about $X_i$ under the surprisal model is thus:

$$I(X_i; R) = \frac{H(X_i|X_1, ...X_{i-1})}{H(X_i|X_1, ...X_{i-1}) + D_{KL}(P_{X_i|X_1,...X_{i-1}}||P^Y_{X_i|X_1,...X_{i-1}})} gH(X_i|X_1, ...X_{i-1}) \tag{19}$$

$$= \frac{g}{1+\alpha} H(X_i|X_1, ...X_{i-1}), \tag{20}$$

assuming that $D_{KL}(P_{X_i|X_1,...X_{i-1}}||P^Y_{X_i|X_1,...X_{i-1}}) = \alpha H(X_i|X_1, ...X_{i-1})$.

**Derivation of surprisal-weighted sampling** We make the same assumptions as the CRUISE model in surprisal-weighted sampling. The only difference is that the number of samples available to store a part of the story is proportional to its surprisal instead of its duration, giving:

$$I(X_i; R) = \frac{g}{1+\alpha} \sum_j \frac{I(X_i; X_j)}{H(X_j)} H(X_j | X_1, ...X_{j-1}). \tag{21}$$

**Fine-tuning on Llama3.2-3b-inst** To test the effect of prior exposure on the mutual information estimates produced by language models, we fine-tuned Llama3.2-3b-inst on transcripts of 301 stories from the Moth Radio Hour. Given that the transcription of *the Moth Radio Hour* was done in-house, it is relatively unlikely that the language models used were trained on these transcripts. We fine-tuned on 3 A100 GPUs with a batch size of 4 for 5 epochs. We adopted an initial learning rate of 5e-6 with cosine decay, and a weight decay of 1e-4. Training loss decreased from 2.7947 to 2.6518. Results suggested that fine-tuning did not lead to large changes in the symmetry and non-negativity metrics that were used to evaluate the mutual information estimates. Results from the fine-tuned Llama3.2-3b-inst model replicated the analyses presented in the main text, supporting the uniform incremental sampling hypothesis (see Fig. S4,S8)).

#### 5.2 Supplemental Results

**Evaluating mutual information estimates** Given that mutual information estimated using different LLMs, concatenation orders, and concatenation methods were not identical, we independently evaluated two theoretical properties of mutual information to identify the optimal combination of LLM and method of estimation (see Methods: Estimating the mutual information of recall and a part of the story $I(R; X_i)$). The first metric evaluates the symmetry of mutual information estimates, operationalized as the root mean square of $I(X_i; R)$ estimated by presenting recall first versus recall last, averaged across all pairs of story parts $X_i$ and recalls (Fig. S2a). The second metric evaluates the non-negative property, operationalized as the proportion of negative $I(X_i; R)$ estimates averaged across all pairs of story parts $X_i$ and recalls (Fig. S2b). For both metrics, lower values indicate more valid estimates. Because we needed an instruction-tuned model to generate recalls for our sampling rate analyses, we selected the instruction-tuned model and concatenation method with the best symmetry score (lowest root-mean square), and then chose the concatenation order with the lowest proportion of negative estimates (Llama3-8b-inst, directly concatenating recall text then story text) for further analysis. Without considering the ability to generate recalls, the LLM-estimation pair with the best symmetry and non-negativity scores was Llama3-8b base model, while presenting recall text then story text using instruction prompting. We reported results of Llama3-8b and others in Fig. S4, S8).

**Validating LLM-derived mutual information estimates using human annotations** We validated our estimates of how much information recalls captured about the story using human annotations. All three stories with human annotations showed high correlations between $I(X; R)$ estimated by Llama3-8b-inst and the number of correct details recalled coded by human raters (Fig. S3, $r > 0.81, p < 1 \times 10^{-10}$). Moreover, $I(X; R)$ significantly predicted the number of correct details even after accounting for the length of recall tokens, $p < 0.001$. We also examined the correspondence between the mutual information estimates and the human annotations at the event level. Results showed high correlations between how much participants' recalls captured about each event of the story, $I(X_i; R)$, and the number of correct details recalled for each event (Story 1: $r = 0.73$, Story 2: $r = 0.60$, Story 3: $r = 0.63$, all $p < 1 \times 10^{-10}$). $I(X_i; R)$ also significantly predicted the number of correct details recalled in each event after accounting for the number of tokens in each event, $p = 0$.

**CRUISE (number of tokens)** While we took $l_j$ (the length of part $j$) as duration in seconds for CRUISE in the main text, we further tested an alternative formulation of CRUISE, treating $l_j$ as the number of tokens tokenized by LLMs. CRUISE (number of tokens) explained similar amount of variance in participants' recalls as CRUISE (duration) (Fig. S4,S5,S8,S9) but they differed in the event boundary analysis. When we divided the story into windows of approximately even number of tokens (Fig. 3a), CRUISE (number of tokens) explained 28.2% of the variance in $I(X_i; R)$, lower than CRUISE (duration) ($Adj.R^2 = 0.298$) but higher than surprisal-weighted sampling ($Adj.R^2 = 0.268$). However, unlike CRUISE (duration), it did not predict that boundary windows are significantly better remembered than inner windows ($p > 0.3$). We also explicitly tested whether event boundary information explains additional variance of participants' recalls beyond CRUISE (number of tokens). When the intercept of window type (boundary vs. inner) was added to the regression, it was significant ($p = 0.037$), suggesting that boundary information captures variance of about participants' memory that is not explained by CRUISE (number of tokens). When only the interaction between window type and CRUISE (number of tokens) was added to the regression, the interaction was not significant ($p = 0.078$), suggesting that the rate of sampling does not differ for boundary and inner windows. When both the intercept and interaction were added to the regression models, neither was significant ($p > 0.1$). These results suggested that while accounting for the shared information structure in the story is important for predicting the memory at event boundaries, the lower speech rate, captured by CRUISE (duration) but not CRUISE (number of tokens), is also a crucial factor.

**Ruling out the shared information confound** One possible reason that CRUISE and surprisal-weighted sampling models both explained significant variance in $I(X_i; R)$ is that certain parts of the story could be more central to the space of all text in the English language, making them share more information with every other part of the story and the recalls, driving up both $I(X_i; R)$ and $I(X_i; X_j)$. To rule out this possibility, we evaluated the ability of CRUISE and surprisal-weighted sampling to predict randomly sampled recalls of non-corresponding stories in our dataset. If the performance of CRUISE and surprisal-weighted sampling is not solely due to certain parts of the story sharing more information with all text in English, then CRUISE and surprisal-weighted sampling should poorly predict the amount of information each part of the story shares with random recalls, $I(X_i; R_{random})$. Practically, for each story, we randomly sampled from recalls of all other 7 stories without replacement for 500 iterations. The number of recalls we randomly sampled equals the number of actual recalls for that story. In each iteration, we calculated the average $I(X_i; R_{random})$ across all random recalls. Each $X_i$ was obtained by dividing the story into windows of approximately even duration, as in Fig. 2. Results showed that CRUISE and surprisal-weighted sampling explain little variance of $I(X_i; R_{random})$ (Fig. S6). For Llama3-8b-inst, the amount of variance CRUISE and surprisal-weighted sampling explained about actual recalls was more than twice the amount of variance they explain about random recalls. These results suggested that CRUISE and surprisal-weighted sampling do not predict a part of the story as better remembered simply because that part of more central to the space of all text.

**Testing the uniform incremental sampling model using consensus events** To test whether the performance of CRUISE and surprisal-weighted sampling are robust to how the story was divided into windows, here we adopted windows of text as consensus events (Fig. S7, S8, S9), instead of windows of even duration as in the main text (Fig. 2). Results suggested that CRUISE (duration), CRUISE (number of tokens), and surprisal-weighted sampling explain more variance about participants' memory of events than surprisal and controls such as total information, duration, and number of tokens. However, because participants' memory of events were strongly influenced by how long the events are (Duration: $adj.R^2 = 0.727$), the numerical differences between the $adj.R^2$ of CRUISE, surprisal-weighted sampling versus controls are smaller than those of the even-duration division we showed in the main text. Still, using a two-level binomial test, CRUISE (duration), CRUISE (number of tokens), and surprisal-weighted sampling predicted participants' memory better than surprisal and controls (Fig. S7e). However, CRUISE did not outperform surprisal-weighted sampling when predicting participants' memory of events, while CRUISE significantly better predicted participants' memory of even-duration windows. These differences could be due to mismatches in the duration and surprisal of certain events that are outliers in event length. Overall, results using consensus events are consistent with those when dividing the windows into even duration, in which CRUISE and surprisal-weighted sampling better predict participants' memory than surprisal and controls.

**Testing the performance of uniform incremental sampling models using other LLMs** Because different LLMs yield different information estimates, we also tested how well CRUISE, surprisal-weighted sampling, surprisal and other controls predict participants' memory using five other LLMs besides Llama3-8b-inst(Fig. S4,S8). Except for results in which mutual information was estimated with Mistral-7b-inst by concatenating recalls and the event text using instruction prompting (Fig. S8b), results across all other LLMs and estimation methods suggested that CRUISE and surprisal-weighted sampling explain more variance in participants' memory compared to surprisal and other controls. One should note that mutual information estimates derived using Mistral-7b-inst with instruction prompting has the worst symmetry among all LLMs (Fig. S2a, mean root mean square of recall last - recall first estimates of Mistral-7b-inst: 88.52 bits, while our best instruction-tuned model Llama3-8b-inst only has a root mean square of 41.92 bits), suggesting that results with Mistral-7b-inst are less valid than those from other LLMs.

**Alternative event boundary analysis** To further validate the information properties and participants' memory at event boundaries, we also split the story into windows of approximately equal duration (Fig. S10a) instead of windows of approximately equal number of tokens as in Fig. 3. Again, the number of windows for each story equals 1.5 times the number of events. A window is considered to be a boundary window if it contains at least one event boundary. The multiplying factor 1.5 is chosen to balance the number of boundary windows and inner windows across eight stories. To avoid abnormally high cross entropy output from LLMs due to syntax violations, the starts and ends of windows were manually adjusted to the closest phrase boundaries. Aligning with the results from the equal-token length splits (Fig. 3), we found that boundary windows have lower speech rate (Fig. S10b, $F = 18.02, p = 3.265 \times 10^{-5}$) and information rate (Fig. S10c, $F = 4.43, p = 0.036$) than inner windows. However, because we explicitly controlled for the duration of windows (Fig. S10e) in this analysis instead of the number of tokens, inner windows contain more tokens (Fig. S10h, $F = 22.42, p = 4.00 \times 10^{-6}$) and higher surprisal (Fig. S10d, $F = 5.18, p = 0.024$) than boundary windows. Given that there is more information to be remembered in inner windows than boundary windows, we found that boundary windows are not better remembered than inner windows (Fig. S10f, $F = 0.43, p = 0.51$), which is predicted by the CRUISE (duration) model (Fig. S10g, $F = 2.71, p = 0.10$). To conclude, when we controlled for the duration of windows, given the lower speech rate and information rate at boundaries, boundary windows are less surprising and are not better remembered than inner windows. When we instead controlled for the number of tokens in windows (see main Results, Fig. 3), boundary and inner windows do not differ in surprisal, and boundary windows are better remembered than inner windows. In either case, memory at boundaries is predicted by CRUISE (duration), which takes into account the lower speech rate and higher shared information at event boundaries. Thus, even though participants' memory at event boundaries seems different when dividing the story into windows of even duration (the current analysis) or even number of tokens (main Results, Fig. 3), these differences are driven by the lower speech rate at boundaries, suggesting the results are compatible.

**Comparing surprisal around event boundaries to baseline** To test whether event boundaries have higher surprisal than non-boundaries, we compared the surprisal of windows centered around boundaries and the surprisal of a null distribution created by randomly sampling windows from the story (Fig. S11). For each story, we took twice the token length of the shortest consensus event as the window size to calculate surprisal. Results suggested that few event boundaries are significantly more surprising than chance. For Story 1 and Story 2, no event boundaries have a surprisal above the 95% of the null distribution. The rest of the stories have maximally 3 boundaries that have a surprisal above 95% of the null distribution: Story 3: 1 out of 27 boundaries; Story 4: 1 out of 17 boundaries; Story 5: 3 out of 20 boundaries; Story 6: 1 out of 16 boundaries; Story 7: 2 out of 24 boundaries, Story 8: 2 out of 11 boundaries. These results suggested that event boundaries are not moments of high surprise, counter to event segmentation theory.

**Information properties of event boundaries at the word level** To better examine how surprisal, speech rate and information rate change as each word is uttered at event boundaries, we also calculated these measures at the word level (Fig. S12) instead of at the time level. Results at the word level aligned with those at the time level, in which surprisal remains mostly constant, while speech rate and information rate decrease at event boundaries.

**Human ratings of the level of gist versus detail in recalls** To measure the level of gist versus detail, human raters parsed the recall transcripts into individual units using an adapted version of Autobiographical Memory Interview coding scheme [66] (see Methods: Recall coding). Each correctly recalled unit was then rated on the scale of 1 to 3, indicating how detailed (3) or gist-like (1) the unit was being recalled. These ratings were then averaged across all units in a recall to yield a mean level of detail score for each recall. Across all three stories with human ratings, recalls rated as more gist-like significantly correlated with lower rate and higher distortion (Fig. S13,S14a,b), supporting our uniform sampling model that information shared across the story (gist) is effectively over-sampled. Moreover, the level of detail ratings also significantly correlated with the proxy of sampling rate in our uniform sampling model (Fig. S14c), the mean attention entropy of the strongest induction head in LLMs from recall tokens to story tokens (see Methods: Evaluation of the attention entropy of induction heads). Higher attention entropy (lower sampling rate) corresponded to more gist-like recalls. These results supported our uniform sampling model that sampling rate correlate with the variation in the level of gist versus detail, and provides an easy-to-compute measure of gist for future studies of narrative recall.

**Sampling rate correlates with the rate-distortion tradeoff** To examine the relationship between sampling rate in our uniform sampling model and the rate-distortion tradeoff, we modeled sampling rate using the attention entropy of the strongest induction head in LLMs (see Results: Rate of sampling predicts the variation of gist versus detail and Methods: Evaluation of the attention entropy of induction heads). Results showed that, across all eight stories, mean attention entropy from recall to stories significantly correlated negatively with rate (Fig. S15a) and positively with distortion (Fig. S15b), suggesting that sampling rate tracks the rate-distortion tradeoff.

**Comparing model-generated recalls to human recalls** While one could observe that model-generated recalls and human recalls overlap on the rate-distortion plots (Fig. 5d), here we explicitly compared the amount of information recalls explained about the story (rate) for model-generated recalls and human recalls (Fig. S16). Results showed that, for every story except for Story 6, attention temperatures exist for which model-generated recalls matched the rate of human recalls using the same generation prompt. Without adjusting for multiple comparisons, rate of model-generated recalls for Story 1 was not significantly different from human recalls at attention temperatures 0.0002 and 0.0003; Story 2 at 0.0001 and 0.0002; Story 3 at 5e-05, 7e-05; Story 4 at 5e-05, 7e-05, 0.0001; Story 5 at 5e-05, 7e-05, 0.0001; Story 7 at 7e-05, 0.0001; Story 8 at 5e-05, 7e-05, 0.0001. For Story 6, because we did not find an attention temperature that matched the rate of human recalls using the current prompt, we prompted the LLM to recall in more detail (See Methods: Recall generation in language models). Using the alternative prompt, without adjusting for multiple comparisons, the rate of model-generated recalls was not significantly different from that of human recalls at attention temperatures 0, 1e-05, 5e-05. These results suggested that LLMs could generate human-like recalls.

#### 5.3 Supplemental Figures

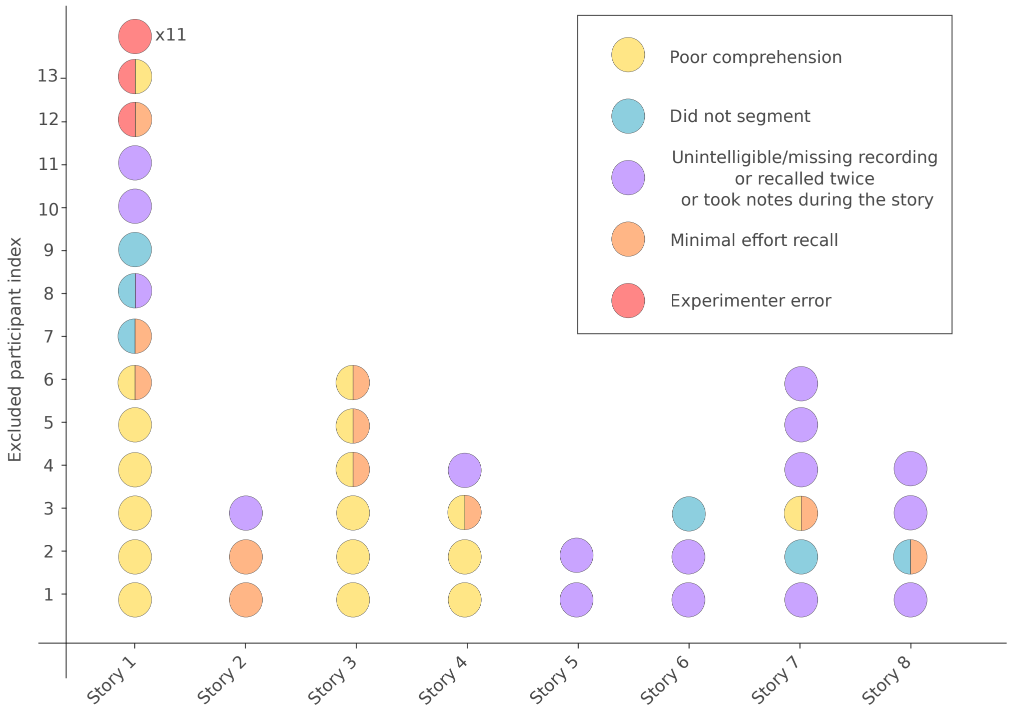


Figure S1: **Breakdown of participant exclusion by story.** Participants were excluded from the calculation of consensus event boundaries if they failed the comprehension check, or did not segment during story listening. Participants were excluded from the recall analysis if they encountered technical difficulties during the recording, or if their recordings were unintelligible, or if they reported recalling twice due to forgetting to press the "start recording" button, or took notes during the story, or spent minimal effort during recall (recalled little to no information about the content of the story) (see Methods: Participants). An additional 13 participants were excluded from the segmentation analysis of Story 1 due to an experimenter programming error, where the comprehension task was incorrectly placed before the recall task, potentially biasing the content of recall.

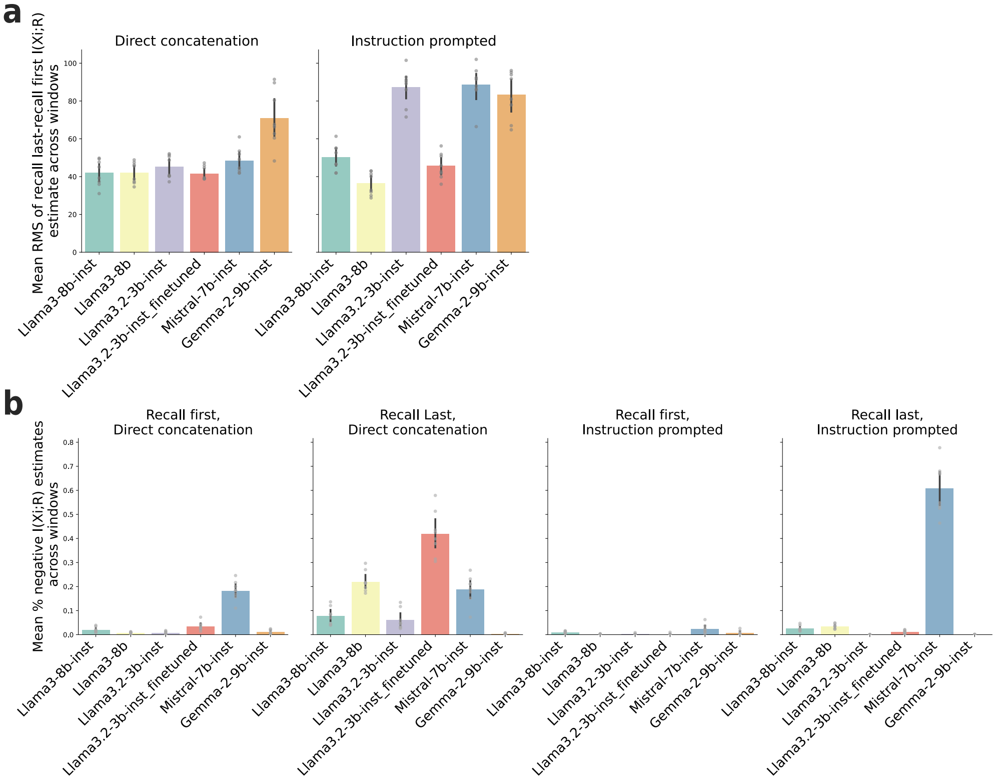


Figure S2: **Metrics for evaluating estimates of $I(X_i; R)$ for each language model. a.** Root mean square of recall last - recall first $I(X_i; R)$ estimates averaged across recalls of all participants and all windows in a story. Windows are derived in two ways: from the consensus event segmentation and splitting the story evenly by duration, in which the number of windows equal to the number of consensus events. **b.** Mean proportion of negative $I(X_i; R)$ estimates across all recalls and all windows in each story, averaged across windows derived from the consensus event segmentation and splitting the story evenly by duration. For both metrics, lower values indicate better estimates.

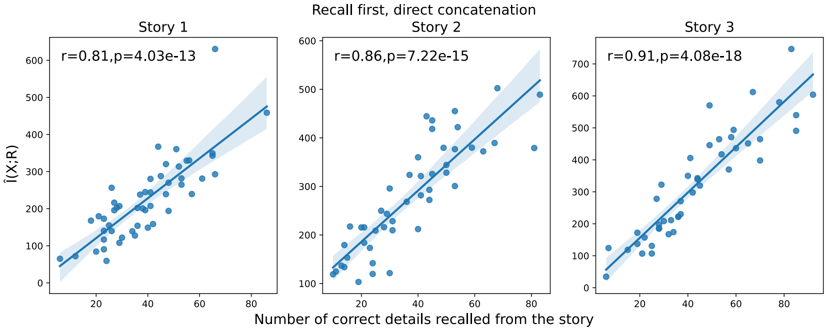


Figure S3: **Correlation between $I(Story; R)$ estimated using Llama3-8b-inst and the number of correct details recalled annotated by human raters.** $I(Story; R)$ was estimated by directly concatenating the recall text then the story text. $I(Story; R)$ estimated by the LLM significantly predicted the number of correct details after accounting for the length of recall tokens, $p < 0.001$.

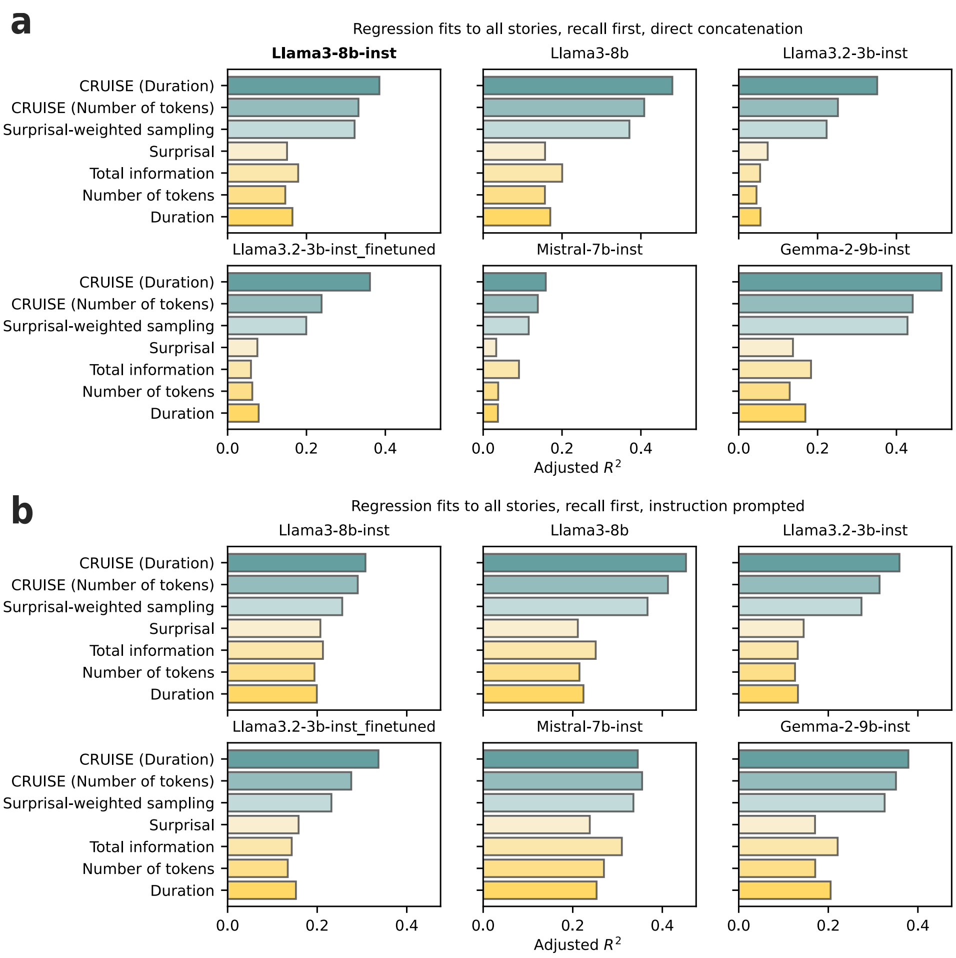


Figure S4: **Comparing the performance of uniform incremental sampling predictors and controls at predicting $I(X_i; R)$ for all stories, across six language models.** Each $X_i$ was obtained by dividing the story into approximately even duration windows. Each linear model contained an uniform incremental sampling predictor or a control predictor, while the slopes of the stories were allowed to vary. $I(X_i; R)$ was estimated by **a.** concatenating the recall directly before the story window and **b.** presenting the recall text and then the story window using instruction prompting. Regardless of the LLM used, CRUISE explained more variance in participants' memory than surprisal or control models.

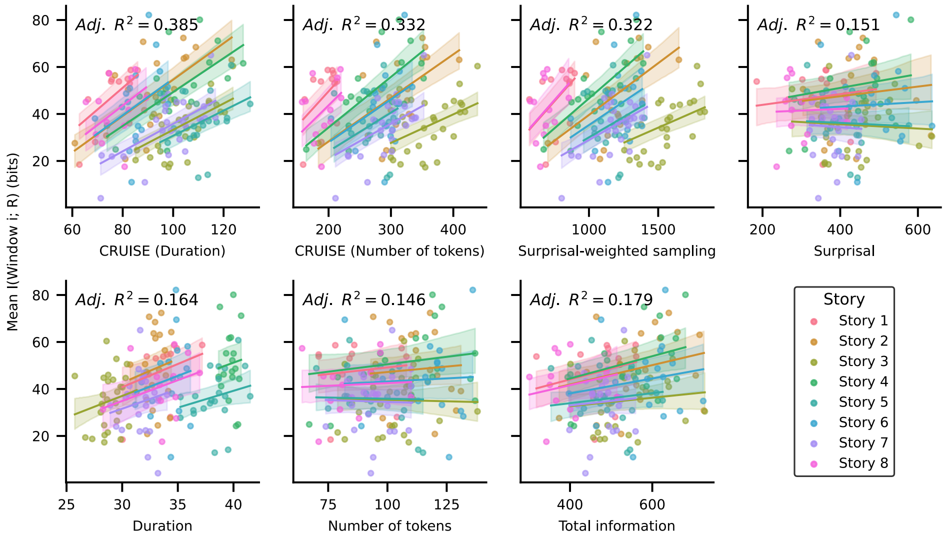


Figure S5: **Predicting $I(X_i; R)$ using uniform incremental sampling and controls.** Each window $X_i$ was created by dividing the story evenly by duration, and adjusting for syntax boundaries. $I(X_i; R)$ was estimated by concatenating the recall directly before the text of the story window using Llama3-8b-inst.

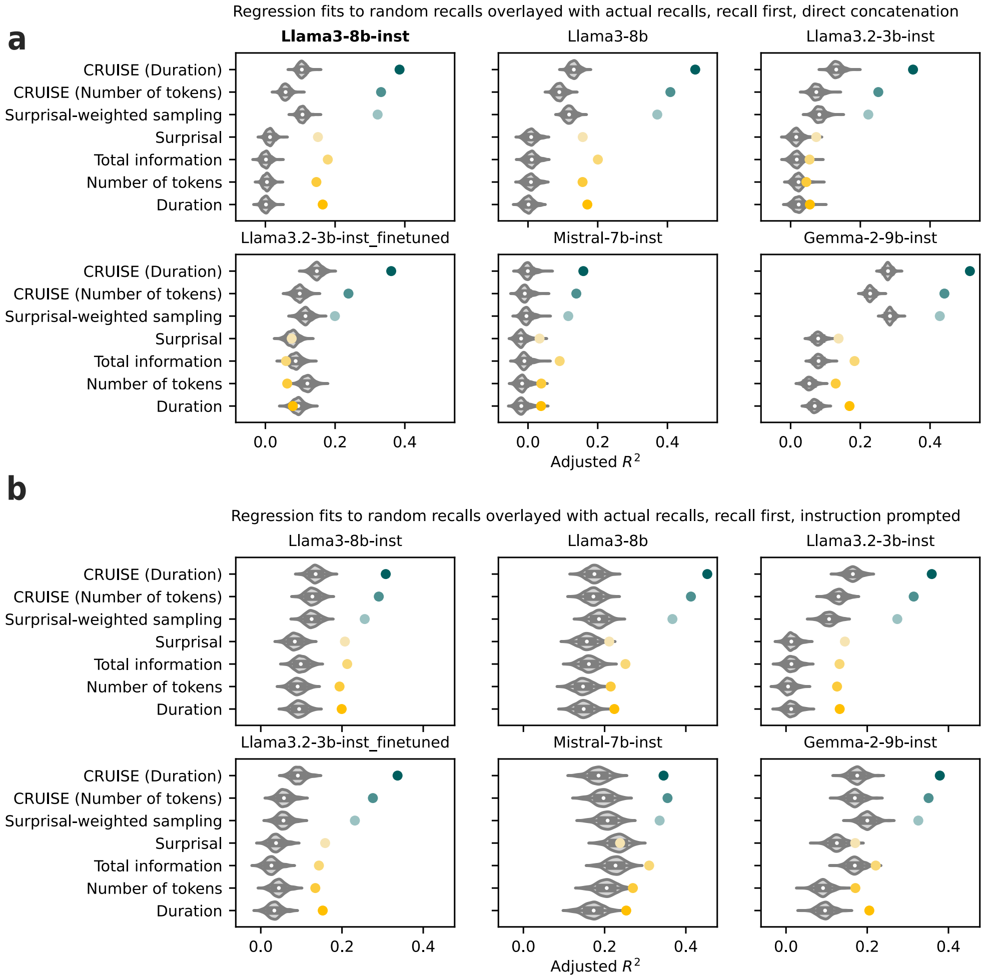


Figure S6: **CRUISE and surprisal-weighted sampling did not predict the mutual information between a part of the story and randomly sampled recalls from other stories.** To rule out the possibility that CRUISE and surprisal-weighted sampling predict participants' recalls because some parts of the stories share more information with all text in English, we tested whether these models could predict random recalls. We randomly sampled recalls from other stories and estimated the mean $I(X_i; R_{random})$, in which each $X_i$ is obtained by dividing the story into windows of approximately equal duration. The gray distributions show the adjusted $R^2$ values from 500 permutations. Inner boxes indicate the interquartile range, and white dots indicate the median. The colored dots indicate the performance of each model in predicting actual recalls of the story. $I(X_i; R_{random})$ was estimated by **a.** concatenating the recall directly before the story window and **b.** presenting the recall text and then the story window using instruction prompting.

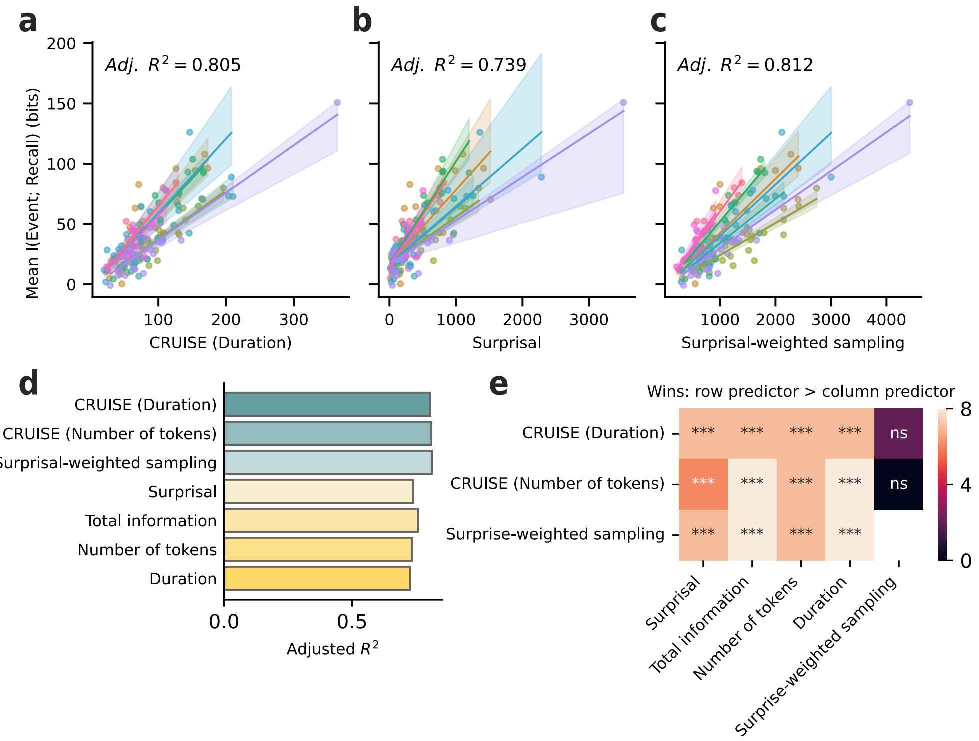


Figure S7: **Testing the uniform incremental sampling model using consensus events. a.-c.** Predicting the mean amount of information participants recalled about each event using **a.** CRUISE, taking the length of each event as its duration in seconds; **b.** surprisal, **c.** surprisal-weighted sampling. Shaded areas indicate 95% confidence intervals. **d.** $R^2$ of predicting the mean amount of information participants recalled about each event using linear regression. Each CRUISE, surprisal-weighted sampling, and control predictor was fitted in a separate linear regression model. Each story was allowed to have its own slope. Because participants' memory of events are strongly influenced by how long the events are, these results are a compressed version of the even-duration division we showed in the main text. **e.** The number of stories (out of 8) in which a row predictor significantly better predicts more participants' recalls than a column predictor. Stars indicate the significance of the second-level binomial test. Results suggested that both versions of CRUISE and surprisal-weighted sampling significantly better predict recalls of more participants than surprisal and controls. However, CRUISE did not outperform surprisal-weighted sampling when the story was divided into consensus events.

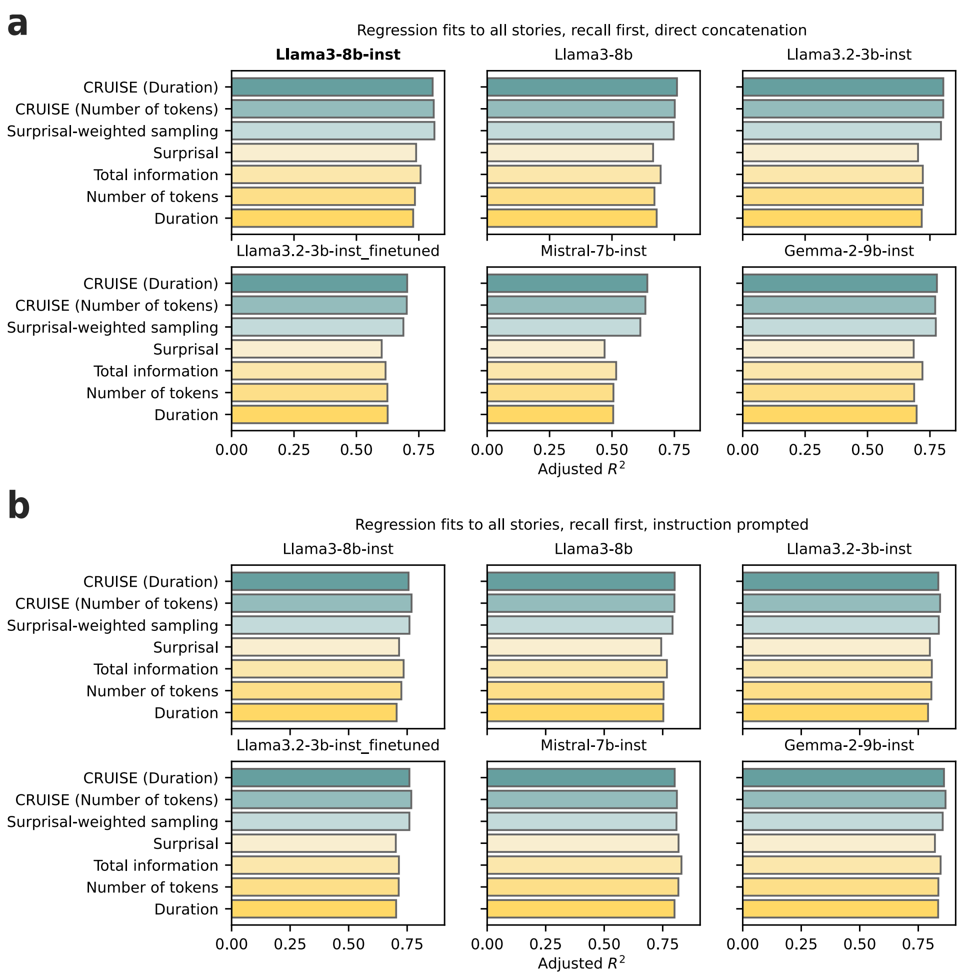


Figure S8: **Comparing the performance of uniform incremental sampling predictors and controls at predicting $I(Event_i; R)$ for all stories, across six language models.** Each linear model contains a uniform incremental sampling predictor or a control predictor, while the slopes of the stories are allowed to vary. $I(Event_i; R)$ was estimated by **a.** concatenating the recall directly before the event text and **b.** presenting the recall and then the event text using instruction prompting. Because participants' memory of events are strongly influenced by how long the events are, these results are a compressed version of the even-duration division we showed in the main text.

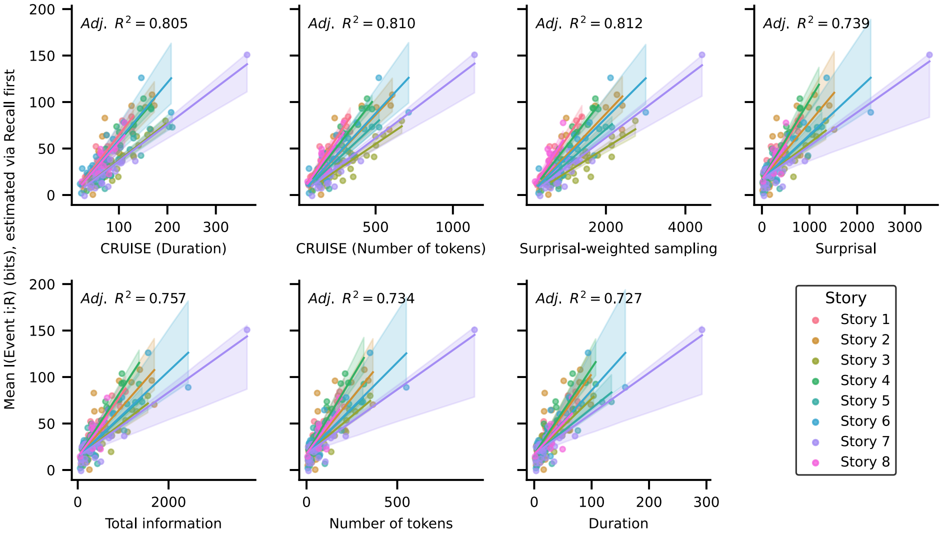


Figure S9: **Predicting $I(Event_i; R)$ using uniform incremental sampling and controls.** $I(Event_i; R)$ was estimated by concatenating the recall directly before the event text using Llama3-8b-inst. Because participants' memory of events are strongly influenced by how long the events are, these results are a compressed version of the even-duration division we showed in the main text.

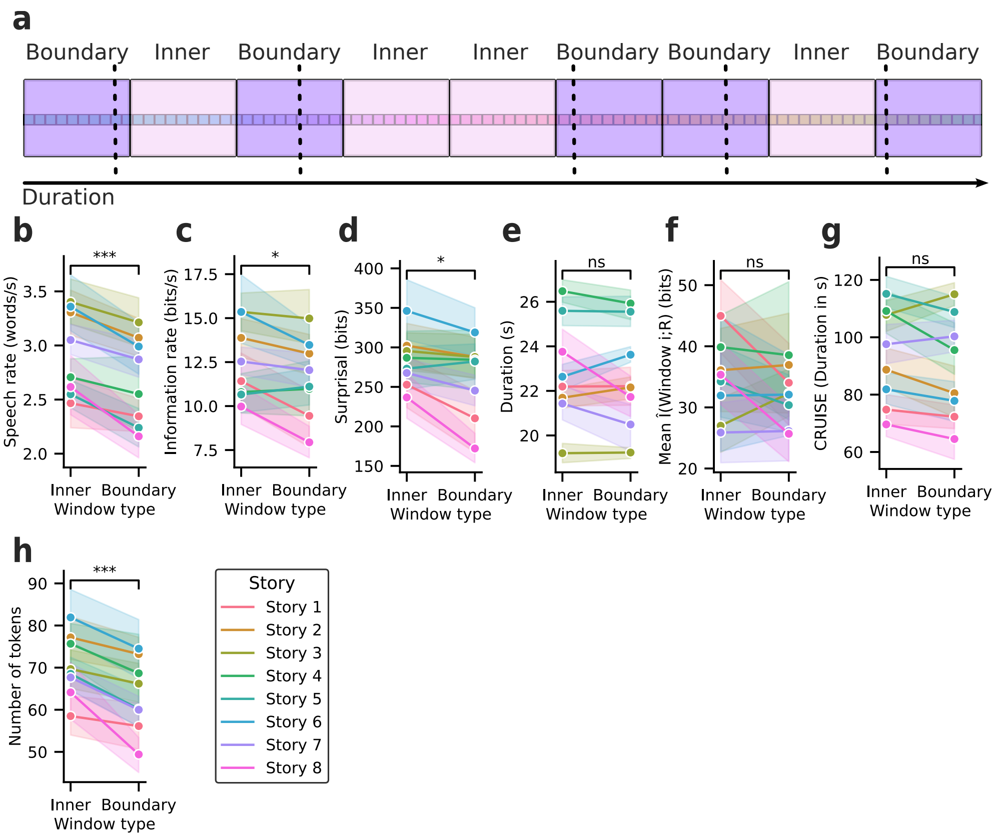


Figure S10: **Alternative event boundary analysis.** **a.** Here we split the story into windows of equal duration, instead of windows of equal number of tokens as in Fig. 3. The number of windows for each story equals 1.5 times the number of events. A window is considered to be a boundary window if it contains at least one event boundary. The multiplying factor 1.5 was chosen to balance the number of boundary windows and inner windows across eight stories. To avoid abnormally high cross entropy estimates from the LLM due to syntax violations, the starts and ends of windows were manually adjusted to the nearest phrase boundaries. **b.** Boundary windows have significantly lower speech rate ($p = 3.27 \times 10^{-}5$) **c.** lower information rate, ($p = 0.036$). **d.** and lower surprisal ($p = 0.024$) than inner windows (shaded area indicated 95% confidence intervals). **e.** Boundary windows and inner windows do not significantly differ in terms of duration ($p = 0.26$). **f.** Boundary windows are not better recalled than inner windows ($p = 0.51$), potentially because there is more information to be recalled in inner windows. **g.** CRUISE (duration) predicts that boundary windows are not better recalled than inner windows ($p = 0.10$). **h.** Boundary windows have significantly more tokens than inner windows ($p = 4.00 \times 10^{-}6$). These results are compatible with those in Fig. 3.

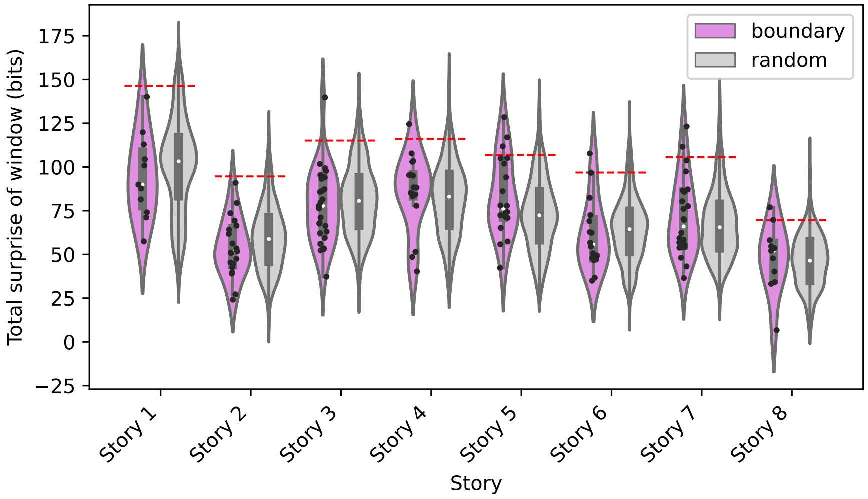


Figure S11: **Surprisal around event boundaries versus baseline.** To test whether boundaries have higher surprisal than non-boundaries, we compared the surprisal of windows around boundaries and the surprisal of a null distribution created by randomly sampling windows from the story. For each story, we took twice the token length of the shortest consensus event as the window size to calculate surprisal. The pink distributions indicate the surprisal of windows centered around event boundaries (black dots represent individual windows). Grey distributions indicate the surprisal of windows of the same size, centered around 1000 randomly selected positions in the story (sampled with replacement). Inner boxes indicate the interquartile range of the null distributions, and whiskers extend to 1.5 times the interquartile range. White dots indicate the median. Red lines indicate the 95% percentile of the null distribution. For Story 1 and Story 2, no event boundaries have a surprisal above the 95% of the null distribution. The rest of the stories have maximally 3 boundaries that have a surprisal above 95% of the null distribution: Story 3: 1 out of 27 boundaries; Story 4: 1 out of 17 boundaries; Story 5: 3 out of 20 boundaries; Story 6: 1 out of 16 boundaries; Story 7: 2 out of 24 boundaries, Story 8: 2 out of 11 boundaries.

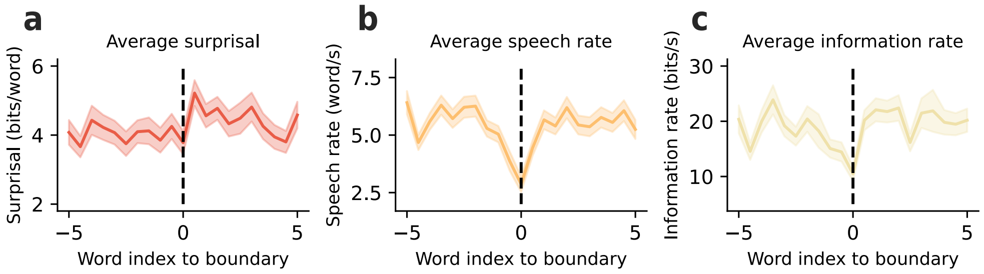


Figure S12: **Average time courses of information properties around event boundaries at the word level.** The average surprisal, speech rate, and information rate from -5 to 5 words around event boundaries. Word index 0 is the last word of the pre-boundary event. The shaded area indicates standard error. This figure is an alternative analysis to Fig. 3j,k,l, in which time courses were calculated for -5 to 5 seconds around boundaries. **a.** Surprisal remains mostly constant around event boundaries. **b.** Speech rate decreases at event boundaries. Here speech rate was calculated as 1/(the start time of next word-start time of current word). **c.** Information rate decreases at event boundaries. Here information rate was calculated as the surprisal of a word/(the start time of next word-start time of current word).

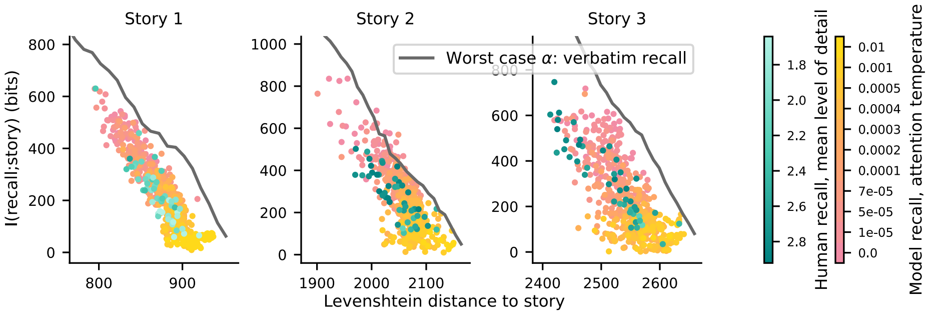


Figure S13: **Rate distortion plot, of individual human recalls and model-generated recalls.** Human recalls (green) are colored by the mean level of detail ratings annotated by human raters, under a coding scheme adapted from Autobiographical Memory Interview. Higher values (darker colors) indicate recalls that are more detailed and less gist-like. Model-generated recalls are colored by the attention temperature used for generation, in which 0 indicate the original model without attention manipulation. Human and model-generated recalls occupy similar levels of rate and distortion. All human recalls and most model-generated recalls lie below the rate-distortion curve of the worst-case $\alpha$, representing someone with no knowledge of the English language, suggesting that both humans and LLM use prior knowledge to efficiently compress the story.

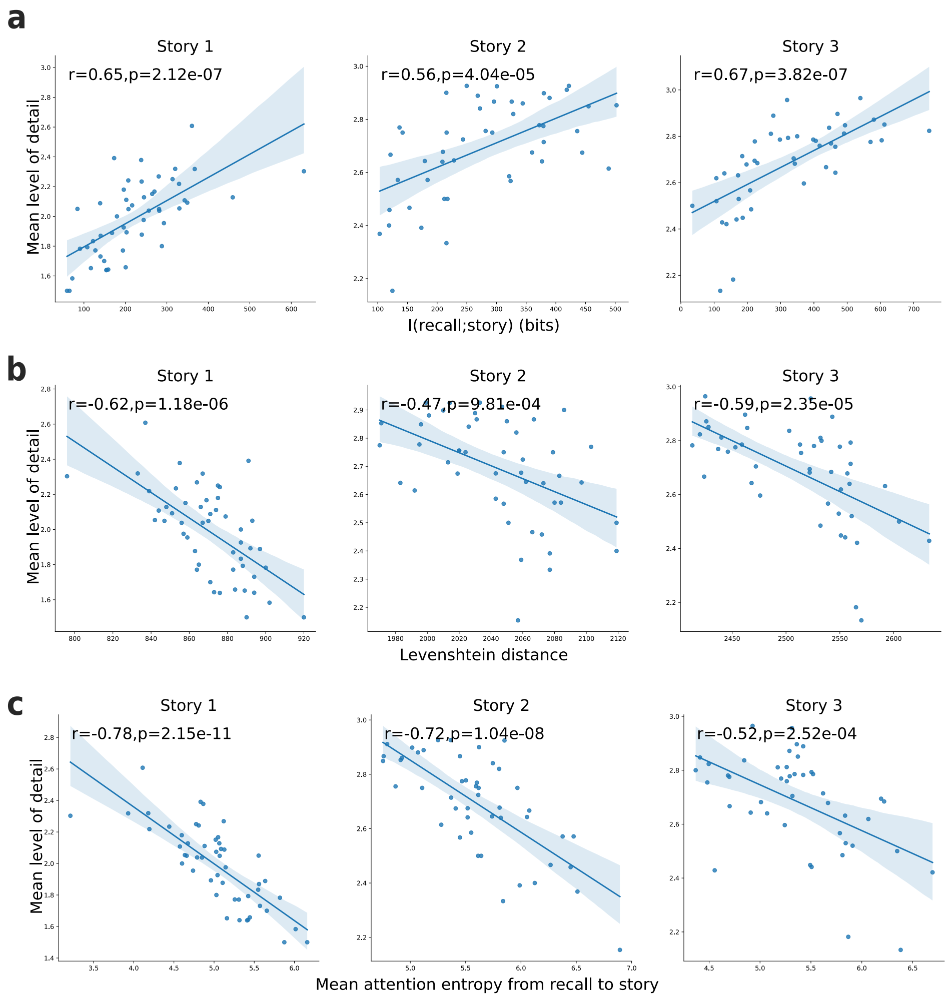


Figure S14: **Correlation of the mean level of detail ratings of human recalls with a.** rate, or the amount of information each recall explains about the story, **b.** distortion, or the word-level Levenshtein distance between each recall and the story, **c.** the proxy of sampling rate in our uniform sampling model, operationalized as the mean attention entropy of the strongest induction head from recall tokens to the story tokens. Results suggested that more detailed recalls rated by a human annotator correlate significantly with higher rate and lower distortion, as predicted by our uniform sampling model. More detailed recalls also correlate significantly with lower attention entropy to the story text, suggesting that sampling rate correlates with the level of detail in one's recall. Results also suggested that attention entropy could act as an easy-to-compute proxy of the level of gist versus details.

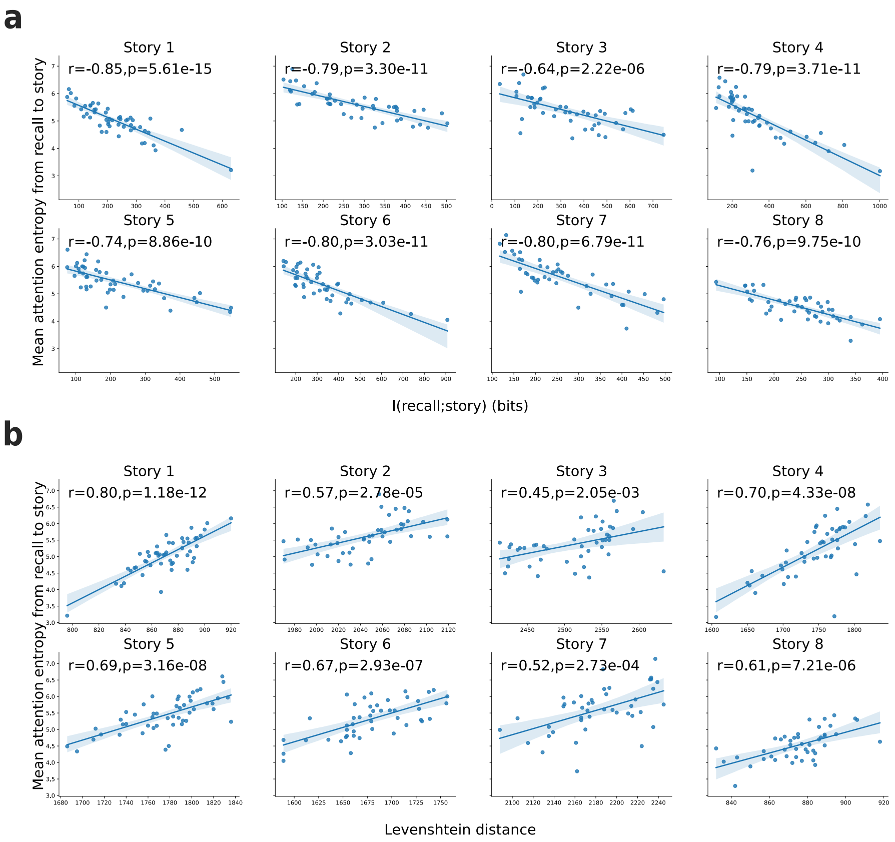


Figure S15: **Correlation of the mean attention entropy of the strongest induction head from recall tokens to story tokens with a.** the amount of information each recall explains about the story. **b.** The word-level Levenshtein distance between each recall and the story. A recall with higher attention entropy to the story text explains less information from the story, and has a larger Levenshtein distance to the story text. Results suggested that attention entropy, the proxy of sampling rate in our uniform sampling model, correlates with the rate-distortion tradeoff.

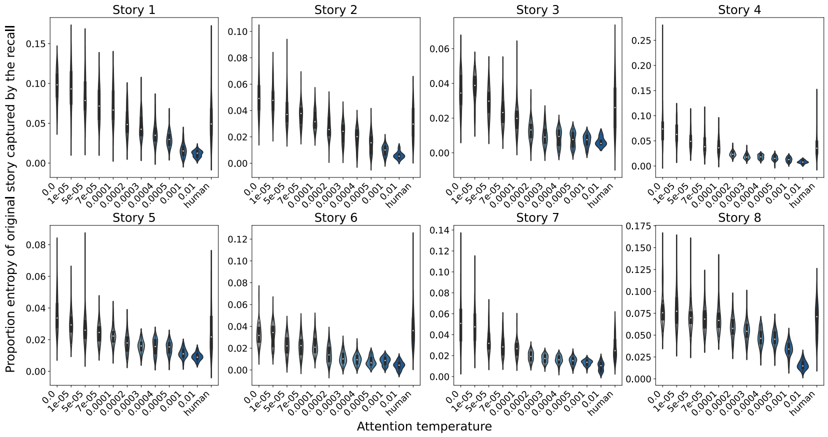


Figure S16: **Comparing the proportion of information recalls explain about the story across model-generated recalls of each attention temperature and human recalls.** Violin plots indicate $I(Story; R)$ of 50 model-generated recalls at each attention temperature, and human recalls. Inner box plots represent the interquartile range. Whiskers indicate 1.5 times the interquartile range, and white dots represent the median. Increasing the attention temperature decreases the proportion of information the generated recalls explain about the story on average, but there is still overlap in the proportion of information explained across attention temperatures, suggesting that the relationship is not deterministic. Using the same generation prompt, in every story except for Story 6, there exist attention temperatures in which model-generated recalls matched human recalls in terms of the amount of information recalls explained about the story. When prompting the LLM to recall in more details, we also found attention temperatures in which the model-generated recalls match human recalls for Story 6.

Table S2: Example model-generated recalls for Story 2: Alternate Ithaca Tom

| scale | transcript |
|---|---|
| 0.00000 | So, there's this guy, and he used to be a custom database application engineer, but he quit his job and bought a Volkswagen van to travel around the country and find his true passion. He didn't really know what he wanted to do, but he wanted something more magical. He drove around for eight months, 25,000 miles, and when he got back to New York, he felt really lost and didn't know what to do. So, he decided to go to a tai chi camp, and that's where he started to feel more connected. He started to feel like he was doing something he loved, and people were really responding to him. He even became the singing asshole of the camp, waking up everyone with his loud singing every morning. But, as he was there, he started to feel like he was living in the wrong life. He kept comparing himself to other people, and he started to feel really unhappy. He kept thinking about this alternate life, where he went to Cornell, studied animal behavior, and became a professor of evolutionary biology. He even had a wife and kids, and they were all really happy. He started to feel really haunted by this alternate life, and he couldn't shake it off. He even went to see a family psychic, who told him that he had a fundamental choice to make in his life. He could have been the hierarchical guy, but he chose to be the piscean, the dreamer, the mystic. The psychic said that if he had chosen to be an academic, he would have been very successful, but his wife would have had an affair with his research partner. But, the guy didn't want to think about that, so he just kept thinking about this alternate life, where he was free and happy. He started to feel like he was stuck in his own life, and he didn't know how to get out of it. He felt like he was just going through the motions, and he didn't know what he wanted to do with his life. That's basically it. I'm not really sure what the point of the story is, but it seems like the guy is just trying to figure out who he is and what he wants to do with his life. He's stuck in this rut, and he doesn't know how to get out of it. |
| 0.00001 | Here's my recall of the story: So, the guy, Tom, used to be a custom database application engineer, but he quit his job when he turned 40 because he felt like it wasn't the right path for him. He bought a Volkswagen van and drove around the country, trying to figure out what he wanted to do with his life. He ended up at a tai chi camp, where he met some interesting people and had a good time. But then he started to feel like he was in the wrong life, and he started to compare himself to others who seemed to have it better. He had this vision of an alternate universe where he went to Cornell, studied animal behavior, and became a professor of evolutionary biology. He even saw a version of himself with a wife and kids who liked to sing. He was really drawn to this alternate life, and it made him feel like he was in the wrong place. Tom started to feel like he needed professional help, so he went to see a family psychic. The psychic told him that he had a choice to make between being an organizational, hierarchical person or a dreamy, artistic person. Tom realized that he had chosen to be the latter, and that if he had chosen the former, he would have been very successful, but also unhappy. The psychic also told Tom that his alternate life was not as perfect as it seemed. His wife in that life was having an affair with his research partner, and he was crushed. Tom started to feel like he was stuck in his current life, and he couldn't shake the feeling that he was in the wrong place. That's my recall of the story! |
| 0.00005 | So, there's this guy, and he used to be a custom database application engineer, but he quit his job because he felt like it wasn't his true calling. He bought a Volkswagen van and drove around the country for eight months, trying to figure out what he wanted to do with his life. He ended up at a tai chi camp, where he started singing and playing music to wake up the other campers. He felt happy and free, but then he started comparing himself to others and feeling like he was in the wrong life. He had this vision of an alternate life, where he went to Cornell and became a professor of evolutionary biology, and he felt like that was the life he was meant to lead. He kept having these visions of alternate universes and alternate versions of himself, and it was really confusing and unsettling. He went to see a family psychic, who told him that he had a choice to make between being a hierarchical, organized person or a creative, artistic person. He realized that he had chosen the latter, and it was causing him a lot of stress and uncertainty. But then the psychic told him that if he had chosen the former path, his wife would have had an affair with his brother or best friend, and he would have been crushed. It was like his alternate life was trying to tell him something, but he didn't know what. Anyway, that's the story. It's all about this guy's journey to figure out who he is and what he wants to do with his life. |

Continued on next page

Table S2: Example model-generated recalls for Story 2: Alternate Ithaca Tom

| scale | transcript |
| --- | --- |
| 0.00010 | So, the story is about this guy who quit his job as a custom database application engineer and bought a Volkswagen van to travel around the country. He wanted to find his real passion and purpose in life. He met a lot of interesting people and had some amazing experiences, but he also felt lost and uncertain about his future. One day, he was driving to a tai chi camp in upstate New York, and he had this vision of an alternate life where he went to Cornell University and became a professor of evolutionary biology. He saw himself with a wife, kids, and a happy life. But then he realized that this life wasn't his, and he felt even more lost and confused. He started to feel like he was stuck in a rut and that he needed to make a change. He went to see a family psychic who did a reading for him and told him that he had a choice to make between being an organized, corporate-type person or a free-spirited, artistic person. The guy was torn between these two paths and felt like he was stuck in the middle. The story is about the guy's journey to find his true path in life and to discover what makes him happy. He's trying to balance his desire for stability and security with his need for freedom and creativity. It's a really relatable story, and I think a lot of people can identify with his struggles and doubts. |
| 0.00020 | Okay, so the story is about this guy who's a bit lost, you know? He quit his job as a custom database application engineer because he felt unfulfilled, and then he bought a Volkswagen van and drove around the country. He ended up at a tai chi camp, where he started to feel more relaxed and happy. But then he started to compare his life to others and felt like he was in the wrong place. He had a vision of an alternate life, where he was a professor of evolutionary biology, and he felt like that was the life he was meant to have. He started to feel like he was stuck in the wrong life. So, he went to see a family psychic, who told him that he had a choice to make between being a corporate guy or a free-spirited artist. He realized that he had chosen the wrong path, and that's why he was unhappy. But then he had another vision, this time of an alternate reality where he was a professor, and he felt like that was the life he was meant to have. He started to feel like he was living in a dream world. I think the story is saying that we all have choices to make in life, and sometimes we have to make sacrifices to follow our true path. But it's also saying that sometimes we can get stuck in our own minds and compare our lives to others, which can make us unhappy. And that's okay, because we can always make changes and try to find our true path. |
| 0.00050 | Hi, so I'm going to try to summarize the story. So, there was this guy who quit his job as a custom database application engineer and bought a Volkswagen van. He drove around the country, and eventually, he ended up at a tai chi camp. He started going there regularly and even became a singer there, singing to wake people up in the morning. But then, he started to feel like he was meant for something more, so he went to a psychic who told him that he was meant to be a professor of evolutionary biology. He became obsessed with this idea and started to feel like he was meant to be someone else. He started to question his life choices and felt like he was stuck in a rut. He even started to feel like he was meant to be someone else, like a different person. He became so obsessed with this idea that he started to feel like he was losing himself. |
| 0.00100 | Here's my attempt to recall the story: So, I was listening to this guy talk about how he was at this camp, and he was talking about how he was feeling really lost and unsure about his life. He said he was trying to find his way and figure out what he wanted to do. He talked about how he used to work in a job that he didn't really like, and how he felt like he was just going through the motions. He said he wanted to find something that made him happy and fulfilled. Then he started talking about how he met this woman, and she was really nice and kind. He said she was very understanding and supportive, and she helped him to see things in a different way. He said she was like a breath of fresh air, and she helped him to feel more alive. He also talked about how he started to pursue his passions and interests, and how that made him feel more happy and fulfilled. He said he started to do things that he loved, and that made him feel more alive. He also said that he started to connect with other people who shared his interests and passions, and that made him feel more connected and supported. I think that's about it. I'm not really sure what else he said, but I think that's the main points of the story. |

Table S3: Example model-generated recalls for Story 3: Ode To Stepfather

| scale | transcript |
|---|---|
| 0.00000 | Okay! So, the story is about this guy, Michael Marquis, who was a big influence in my life when I was a kid. He was like a mentor, a father figure, and a friend all rolled into one. He was a Vietnam vet, a professional athlete, and a ski instructor, and he had this really cool MG car. He was the first person to really tap into my "momma's boy complex" and help me feel like a man. He taught me how to play football, chop wood, and even how to shoot a shotgun. He was always trying to help me become more confident and self-assured. He even took me on adventures, like going to see movies and taking me on dates with other women. Yeah, it was a little weird, but it was also really cool. But, things got weird when my mom started dating other guys, and Michael would take me on these wild adventures with these women. And then, one day, my mom's underwear and lingerie were nailed to the outside of our cabin, and I was like, "What the heck is going on?" Anyway, Michael was a big part of my life until my dad got remarried, and my mom started dating other guys. I got kicked out of a gifted and talented program for telling a kid that his painting sucked, and then I got in a fight and got kicked out of school. But, Michael came back into my life for my birthday party, and it was like one of the greatest days of my life. He took me and my friends on a wild ride in his MG, and we went to Chuck E. Cheese's and saw George Burns. It was like, the ultimate birthday party. But, then he left, and I was devastated. I didn't want him to see me crying, so I told him goodbye, and he promised to stay in touch. And, yeah, he did. He even told me he loved me, and that he was gonna stay in touch with me. And, for a while, he did. But, then he disappeared, and I never saw him again. Years later, my mom got a message from him, saying that he saw some movie I was in and thought I was doing great. And, yeah, it meant a lot to me. Because, even though I never saw him again, he's still been a big influence in my life. He's like, the secret influence that's always been there, guiding me and helping me become the person I am today. |
| 0.00005 | Okay, so... (pauses) Oh boy, this is gonna be tough. (laughs) Alright, so the story is about this guy, Michael Marquis, who was a big influence in my life. He was like a mentor or something. I met him when I was seven, and my mom had brought him home. He was a Vietnam vet, played college football and baseball, and was just a really cool guy. He had this MG car that was super cool, and he taught me how to play football and stuff. Anyway, we moved to Vermont, and he took me under his wing. He taught me how to chop wood, and we'd go on these adventures together. He was like a father figure to me, and I looked up to him a lot. He even took me to see "The Man Who Would Be King" and I cried at the end, and he was like, "Come on, it's just a movie." But then things got weird. My mom would go out with other guys, and Michael would take me on dates with them. It was all kinda strange. And then one day, we moved to Atlanta, and my dad got remarried. I was really upset about that, and I got kicked out of this gifted program at school because I told someone their painting was bad. (laughs) Anyway, Michael showed up at my birthday party, and it was like, the best day ever. He took us on a wild ride in his MG, and we went to Chuck E. Cheese's and saw George Burns. It was just so cool. But then he had to leave, and I was devastated. I didn't want him to see me crying, so I just... (pauses) Yeah, it was a tough time. And then I never saw him again. But my mom got a message from him a few years ago, saying he saw some movie I was in and thought I was doing great. And that was it. He was the secret influence in my life, and I'll always carry him with me. (pauses) That's the story... I think. (laughs) |

Continued on next page

Table S3: Example model-generated recalls for Story 3: Ode To Stepfather

| scale | transcript |
| --- | --- |
| 0.00007 | So, the story is about a guy who had a really influential person in his life named Michael Marquis. He met Michael when he was seven years old, and Michael was like a father figure to him. Michael was a Vietnam vet, a professional athlete, and had a lot of cool stuff like a '33 MG and a German Shepherd. He was really into Texas and the Dallas Cowboys, and he helped the guy overcome his momma's boy complex. Anyway, Michael took the guy and his mom to Vermont, where they lived in a one-room cabin. Michael taught him how to chop wood, and they went on adventures together. The guy loved Michael, but his mom had some issues, and they eventually left Vermont. The guy's dad got remarried, and they moved to Atlanta, Georgia. The guy got into some trouble at school and had to write an essay about why saying someone's painting sucked was bad. He also got into a fight and got kicked out of a gifted program. The guy's mom and Michael remained close, and Michael would visit them sometimes. The guy loved Michael, but he was also really hurt when Michael left. The guy's mom would get upset when Michael came around, and the guy felt like he was stuck between his mom and Michael. Eventually, Michael left the guy's life, but the guy never forgot about him. He carried Michael with him every day and considered him the secret influence of his life. |
| 0.00010 | Okay! So, there was this guy, Michael Marquis, who was like a mentor to me when I was a kid. He was a Vietnam vet, a professional athlete, and just a really cool guy. He lived with my mom for a while, and I was really close to him. He taught me how to play football, and we'd go on adventures together. He was like a father figure to me, but also kind of a bad influence. He'd take me on dates with other women, and my mom would get mad at him. He was also really mean to my mom, and I didn't like that. Anyway, we moved to Vermont, and Michael would take me on these wild adventures. He was always trying to teach me how to be a man, and I guess it worked because I started to feel more confident. But then, things got weird. My mom would get mad at him, and he'd get mad back. It was like they were in this toxic relationship or something. Then, my dad got remarried, and I was really upset about it. I didn't get invited to the wedding, and it felt like my whole world was turned upside down. I got kicked out of this gifted program at school because I told a kid that his painting was bad. My mom was really mad at me, and I felt like I was in trouble all the time. But then, Michael showed up at my birthday party, and it was like a dream come true. He took me and my friends on this wild adventure, and we had the best time. He was like a superhero or something. But then, he had to leave, and I was devastated. I didn't want him to go, but he said he'd stay in touch. After that, I never saw him again, but I've always carried him with me in my heart. He was like a secret influence in my life, and I'll always be grateful for the time we had together. |
| 0.00020 | Okay! So, the story is about this guy, and he's talking about this person, Michael Marquis, who was a big influence in his life. Michael was like a mentor or a role model to him, and he met him when he was just a kid. Michael was a cool guy, he was a Vietnam vet, a professional athlete, and he had a lot of wisdom to share. The guy is saying that Michael was the first person to really understand him, and he was like a father figure to him. Michael taught him how to play football, and they would go on adventures together. The guy loved Michael, but he also felt like he was trying to be too cool for him. There was this one time when the guy's mom was dating Michael, and he felt like he was being left out. He was jealous of the attention Michael was giving his mom, and he felt like he was being replaced. But Michael was still nice to him, and he would take him on adventures and teach him new things. The guy is saying that Michael was the one who made him feel like he was a man, and he loved him for that. But he also felt like Michael was trying to control him, and he didn't like that. He felt like Michael was trying to make him into something he wasn't, and that was hard for him to deal with. Anyway, the guy and Michael had a falling out, and they didn't talk for a long time. But the guy never forgot about Michael, and he still thinks about him to this day. He's saying that Michael was a big influence on his life, and he's grateful for the time they spent together. That's the story, I think! |

Continued on next page

Table S3: Example model-generated recalls for Story 3: Ode To Stepfather

| scale | transcript |
|---|---|
| 0.00050 | Here's my attempt to recall the story: So, I think the story is about this person who has a lot of fond memories of their childhood, especially with this one guy, Michael, who was like a mentor or a role model to them. They mention how Michael would take them on adventures and teach them things, and they seem to have a really strong bond. The person also talks about how they grew up in a household with a lot of chaos and uncertainty, and how Michael was one of the few constants in their life. They mention how Michael would take them on trips and teach them new things, and how they would often get into mischief together. The person also talks about how they have a lot of regrets and "what ifs" in their life, and how they wish they could go back and relive some of those moments with Michael. They seem to have a lot of admiration and respect for Michael, and they're grateful for the impact he had on their life. Overall, the story seems to be about the importance of having positive role models in your life, and how those relationships can shape who you become and what you do with your life. |
| 0.00100 | *ahem* Okay, so I'm gonna try to summarize this story for you. So, the guy was talking about how he used to live with his mom and his stepdad, and they would always fight and argue. He would always try to stay out of the way and avoid getting involved in their fights. One day, his mom told him that she was leaving his stepdad and taking him and his sister with her. He was really upset and didn't want to leave his home and his friends. But his mom told him that they had to leave because it wasn't safe for them to stay there anymore. |
